# Quantitative Fluorescence Imaging with SI-Traceable Radiance: Radiometric Fluorescence Characterization with a Calibrated Solid-State Emitter

**DOI:** 10.1101/2025.10.15.677906

**Authors:** Alberto J. Ruiz, Edwin A. Robledo, Eammon A. Littler

## Abstract

**Significance:** Fluorescence imaging remains largely qualitative and device-specific, limiting reproducibility and intersystem comparisons. Advancing toward quantitative imaging requires a radiometric characterization framework that provides SI-traceable units while explicitly addressing the interdependent factors that govern image formation.

**Aim:** Establish and evaluate a radiometric characterization framework that converts device-native counts to SI-traceable imaged radiance (µW·cm^−2^·sr^-1^) and aggregate fluorescence yield (sr^-1^) while accounting for interdependent factors that influence fluorescence image formation.

**Approach:** A calibrated Lambertian solid-state radiometric emitter target (RET) was combined with a three-step radiometric framework consisting of the Radiance Transfer Curve (RTC), the Radiance Imaging Transform (RIT), and the Fluorescence Imaging Transform (FIT). The RTC establishes system responsivity as a function of radiance; the RIT applies this calibration at the pixel level to map digital counts to SI-traceable radiance; and the FIT performs pixel-wise excitation normalization of the RIT image to produce an aggregate fluorescence yield image. The framework was tested through distance and aperture invariance, digital-vs-physical ROI analyses, RTC acquisition, and application of the RIT and FIT to an ICG concentration target and a breast lumpectomy phantom.

**Results:** The RET exhibited Lambertian behavior, with no significant dependence of the measured radiance on distance or aperture; imager responsivity (*R*_*λ*_) also remained invariant within uncertainty across working distances and f-numbers. Digitally masked ROIs reproduced *R*_*λ*_ obtained with matched physical apertures, enabling ROI and pixel-level radiance transfer. RTCs acquired over 49 radiances captured sensor and processing nonlinearities. The RIT provided a per-pixel mapping from counts to radiance (µW·cm^−2^·sr^−1^). Applying pipeline-specific RTCs, the RIT and FIT reconciled large discrepancies across RAW, 8-bit, and log10 image processing pipelines, yielding closely aligned radiance-concentration curves and improved SBR/SNR/CNR agreement. In a breast lumpectomy phantom, FIT produced SI-traceable aggregate fluorescence yield (sr^−1^) images and absolute contrast metrics in an anthropomorphic geometry.

**Conclusions:** The combined framework converts device-native counts into SI-traceable radiance and aggregate fluorescence yield at the pixel level, providing a practical basis for reproducible quantitative fluorescence imaging. The feasibility results across distance/aperture tests, ROI analyses, image pipelines, and phantom imaging indicate readiness for broader evaluation. Future work will establish formal uncertainty budgets and assess robustness across devices, geometries, and excitation conditions to support adoption as a quantitative reporting standard.

## 1 Introduction

Fluorescence imaging and sensing are now widely used in molecular and cellular biology, in preclinical studies, and in standard clinical workflows.^1–3^ In molecular and cellular biology, techniques such as fluorescence microscopy, super-resolution imaging, and lifetime imaging enable detailed investigation of protein dynamics, cellular morphology, and microenvironmental physiology.^4,5^ In preclinical research, whole-animal fluorescence imaging enables longitudinal assessment of probe biodistribution, disease progression, and treatment response in small-animal models,^6,7^ while larger animal models, particularly porcine, are increasingly used for indication development, device performance evaluation, pharmacokinetics validation, and surgical training.^8–10^ Clinically, fluorescence-guided imaging (FGI) and fluorescence-guided surgery (FGS) augment conventional white-light visualization by providing molecular and physiological contrast enabling oncologic margin delineation, perfusion evaluation, lymphatic mapping, and identification of critical anatomy.^11–14^

Although FGI and FGS are increasingly used in the clinical setting, most imaging remains device-specific and qualitative, thereby limiting comparison across devices, sites, and patient cohorts.^15,16^ Current systems typically report device-native digital numbers rather than SI-traceable radiometric units (i.e., quantities traceable to the International System of Units through calibration with reference standards maintained by national metrology institutes), which obscures the relationship between measured signal and true fluorescence emission radiance.^17,18^ This limitation makes it difficult to differentiate the performance of contrast agents from that of the imaging devices.^16^ The lack of radiometric traceability is compounded by the absence of broadly adopted quantitative standards and test methods, leading to substantial variability in device performance observed in translational and clinical devices.^18–20^ Additionally, commonly reported contrast metrics such as tumor-to-background and signal-to-background ratios are highly sensitive to acquisition parameters, ROI selection, and image-processing pipelines, reinforcing the need for radiometric frameworks that yield SI-traceable quantitative fluorescence to support rigorous clinical evaluation and regulatory assessment.^15,16,21^

Quantitative imaging and cross-system comparison are challenged by the many interdependent factors that influence fluorescence image formation, including excitation irradiance (spectrum, intensity, and spatial homogeneity), tissue optical properties (absorption, reduced scattering, and autofluorescence), optical throughput (numerical aperture, filter/dichroic bands), imaging geometry (working distance and angles), sensor characteristics (quantum efficiency, gain, dynamic range, dark and thermal noise), and the image-processing pipeline (gamma curves, denoising, compression, baselining). Without radiometric characterization and standardized reporting (e.g., imaged fluorescence radiance), two imagers can generate different contrast values for the same field, complicating multicenter trials, pooled analyses and meta-analyses, and ultimately limiting the generalizability of clinical results. This variability also raises concerns that different devices may not deliver comparable visualization or clinical outcomes, creating the risk that subsequent systems cannot reproduce prior performance in the absence of truly comparable imaging data.

Previous work on quantitative fluorescence imaging has mostly relied on relative calibration to fluorescence phantoms, lacking true traceability to SI units for the imaged fluorescence.^15,16,18,19,21–26^ Within SI-traceable efforts, two distinct routes are generally recognized, one based on the amount of substance (e.g., equivalent number of reference fluorophores in flow cytometry)^27^ and the other based on optical metrology with calibrated light sources and solid-state detectors.^20,28^ Recent methods, largely led by the National Institute of Standards and Technology (NIST), based on radiance calibration and Lambertian emission, represent the most substantive progress toward true quantitative fluorescence-guided imaging.^20,29–33^ These methods employ calibrated radiance sources and known collection geometry to establish device responsivity and to express captured fluorescence in SI-traceable radiance (W·m^−2^·sr^-1^), with explicit reporting of excitation irradiance at the imaging plane.^31^ Litorja (2022)^29^ formalized radiance transfer from SI-traceable tissue phantoms, introducing the aggregate fluorescence yield factor (emission-radiance-per-excitation-irradiance with units of sr^-1^) and utilizing *étendue* invariance^34^ to support absolute, comparable measurements. Establishing Lambertian (or near-Lambertian) behavior of reference materials is central to this transfer, reducing geometry-dependent bias and enabling distance-invariant mapping of the imaged radiance for fixed optical geometries.^30,31^ Related efforts in SI-traceable quantitative luminescence, particularly the use of solid-state emitters as calibration standards, further support ongoing work toward standardized and reproducible calibration approaches.^35–39^ Together, these advances provide a foundation for expanding SI-traceable measurements to imaged fluorescence for FGI and FGS devices; however, current implementations typically characterize the imager at a single radiance point and implicitly assume linear response, which does not capture sensor non-linearities, photon-noise regimes, or transformations introduced by image-processing pipelines. Accordingly, there remains a need for a comprehensive radiometric framework that characterizes responsivity across the full signal range, enables pixel-level propagation of radiance, and incorporates excitation-normalized fluorescence metrics reported in absolute radiometric units to enable reproducible system characterization and cross-system analysis.

Previous work on non-fluorescent radiometric imaging has been extensive, particularly focused on photon transfer and scene brightness applications.^40–42^ The photon-transfer method calibrates sensors by converting device-native digital numbers (N) to electrons (e^−^) from measured signal–noise statistics, treating the camera as a black box.^40^ The resulting photon transfer curves quantify gain (e^−^/N), read and shot noise, full-well capacity, linearity/nonlinearity, and usable dynamic range in absolute units, enabling subsequent radiometric imaging. This fundamental relationship between N and photoelectrons, governed by the semiconductor bandgap of the photosites and digitization of the resulting charge, has enabled imaged irradiance (W·cm^−2^) to serve as the basis for radiometric calibration and scene brightness recovery across a wide range of applications including astronomy, machine vision, photometric measurements, and high-dynamic range imaging.^41–46^

This work aims to enable reproducible, SI-traceable quantitative fluorescence reporting across FGI and FGS devices. Here, we established radiance transfer using a calibrated solid-state emitter with adjustable radiance output, confirming Lambertian behavior that supports distance-invariant calibration across low- and high-signal regimes. Building on NIST radiance-transfer methods and photon-transfer concepts, we developed the Radiance Transfer Curve (RTC) to capture device responsivity throughout the full system dynamic range within a spectral band, explicitly reflecting non-linear sensor behavior, optical configuration, acquisition parameters, and the effects of image-processing pipelines. We also introduced the Radiance Imaging Transform (RIT), which applies the RTC at the pixel level to map images from digital numbers N to imaged radiance (W·m^−2^·sr^−1^), We additionally define the Fluorescence Imaging Transform (FIT) as a per-pixel excitation normalization that reports the imaged aggregate fluorescence yield (sr^−1^), which enables direct cross-system comparisons by minimizing the impact of instrument-specific response and distance-dependent fluorescence excitation effects. The following sections present the theoretical and mathematical framework for radiance transfer, RTC, RIT, and FIT together with radiometric evaluations under varied imaging parameters and fluorescence-phantom experiments, demonstrating feasibility across a range of conditions.

## 2 Methods

### 2.1 Calibrated Solid-State Emitter: Radiometric emitter target

The calibrated solid-state emitter (radiometric emitter target, RET) is shown in **Fig 1** (QUEL Imaging, RRL-825-WR01-QUEL01) alongside the radiance vs. voltage curve, spectrum, and intensity profile characterizations. The RET consists of an NIR LED, diffuser stack, aluminum base, and 3D printed housing components [**Fig 1(a)**]. It is voltage controlled to produce a radiance output of ∼ 0.1 – 100 µW·cm^−2^·sr^−1^ [**Fig 1(b)**] with a spectral peak output centered at 827 nm with a full-width half-maximum of ∼35 nm. The RET output spectrum overlaps that of ICG in plasma [**Fig 1(c)**], enabling imaging with most systems indicated for use with 800 nm channel fluorophores. Although we used an 820-830 nm NIR band for ICG imaging applications, the calibration methodology is wavelength-agnostic: substituting the LED emitter allows alignment of the RET spectrum with fluorophore-specific emission bands. This target provides a pseudo-flat emission profile [**Fig 1(c,d)**] which facilitates the use of region-of-interest (ROI) analysis. The RET calibrated radiance values were determined using **Equation (1)** from measurements obtained with a NIST-traceable photodiode (Thorlabs Inc., S120VC) positioned behind a 5 mm pinhole aperture (Thorlabs Inc., P5000K) at 100 mm in a custom 3D-printed enclosure. The 1.1-1.34 V driving-voltage range corresponded to a measured radiance emission range of 0.04 – 237 µW·cm^−2^·sr^−1^.

**Fig. 1:**
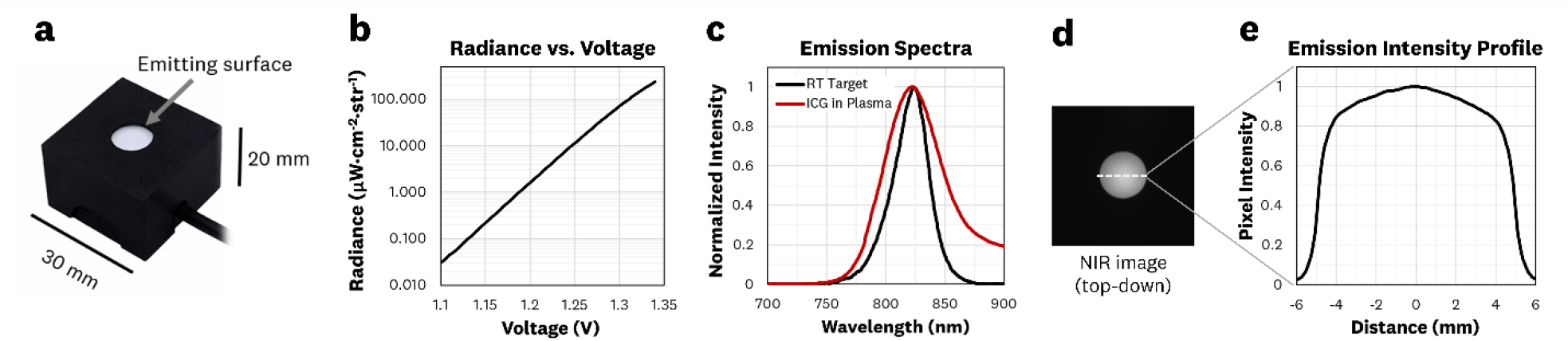
The RET alongside radiance, spectrum and intensity profile characterization: **(a)** The RET has a ∼10 mm emitting output aperture with **(b)** voltage control to produce a radiance output of ∼0.1-100 µW·cm^2^·sr. **(c)** The spectral output of the RET falls within the spectra of ICG in plasma, indicating adequacy for use with most 800 nm channel fluorescence systems. **(d)** The RET has a pseudo-flat emission profile providing a diffuse output for imaging applications.

### 2.2 Evaluating the Lambertian emission of the RET

To enable the conversion of imager response to SI units, the RET must exhibit a Lambertian output.^30^ In brief, Lambertian sources have radiance that remains invariant with distance and solid angle, enabling direct mapping of an SI-traceable radiance (W·m^−2^·sr^−1^) to the imager response.^29,30^ The radiance emitted by a Lambertian source (*L*_*src*_) can be determined by the detected power from a calibrated photodiode power sensor (Φ_*ps*_) for a given emitter source area (*A*_*src*_) and associated solid angle (*Ω*_*src*_). As shown in **Fig 2(a)**, *Ω*_*src*_ is defined by the aperture stop (*A*_*apt*_) and the source-to-aperture distance (*d*). Mathematically the radiance is defined as

**Fig. 2:**
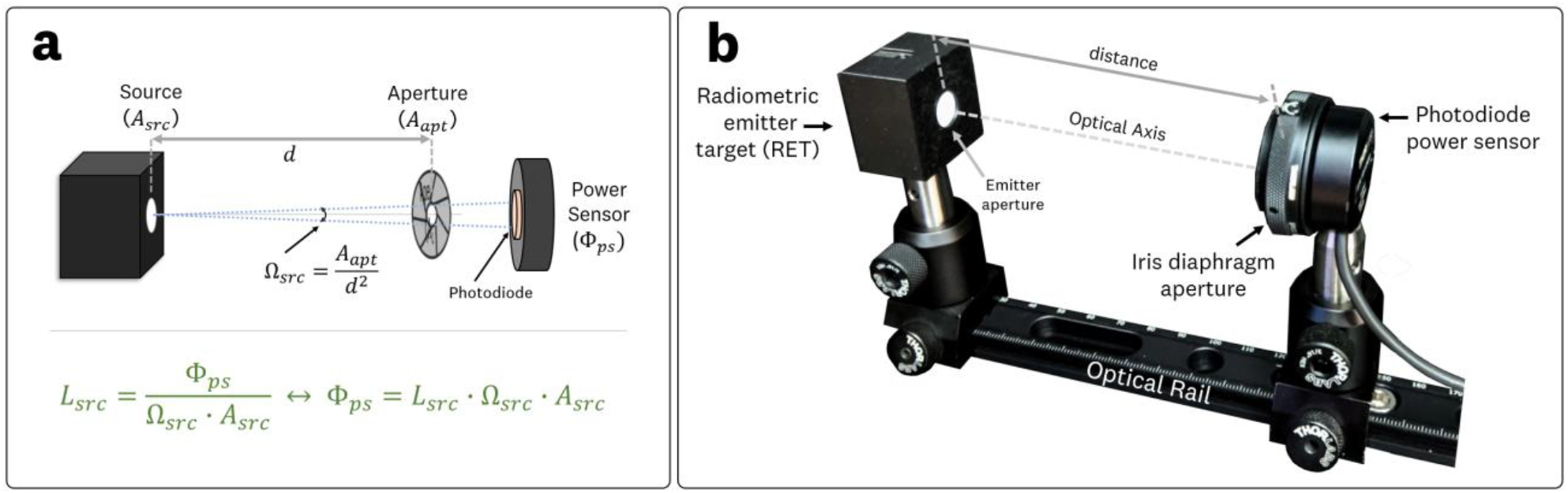
Optical set-up for assessing the RET’s Lambertian behavior: **(a)** Geometric optics schematic for radiance *L*_*src*_ measurement, illustrating the light transmission from source to detector alongside relevant quantities and **(b)** experimental set-up for testing the Lambertian behavior of the RET.

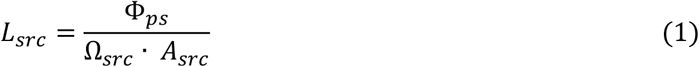

where Ω_*src*_ = *A*_*apt*_/*d*^2^. Conversely, if the radiance is known, the detected radiant flux (i.e. power) is given by

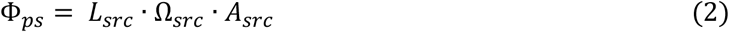

These mathematical relationships indicate that a calibrated photodiode can be used to measure the radiance output of a source, by additionally measuring *d, A*_*src*_, and *A*_*apt*_.

**Fig 2(b)** shows the experimental setup used for assessing the RET’s Lambertian behavior, which utilized a calibrated photodiode (Thorlabs Inc., S120VC) and a variable iris diaphragm aperture (Thorlabs Inc., SM1D12C) mounted on an adjustable optical rail. The RET and photodiode were aligned along the optical axis at a *d* = 400 mm to achieve consistent optical power measurements within the tested 25 - 400 mm range. To test the radiance invariance with distance, the aperture was fixed at 5.0 mm (*A*_*apt*_ =19.6 mm^2^) and the detected power recorded at *d* = 25, 50, 75, 100, 150, 200, 250, 300, 350, and 400 mm. To test the solid-angle invariance, the distance was fixed at *d* = 100 mm, while the aperture diameter varied at 2, 3, 4, 5, 6, 7, and 8 mm utilizing the markings of the variable iris diaphragm aperture. Each measurement series was acquired in triplicate. Furthermore, the true aperture diameter at these diaphragm markings were measured in triplicates using a calibrated caliper (Mitutoyo, 500-196-30), such that the aperture diameters used in the radiance calculation were of 1.86, 2.83, 3.90, 4.93, 5.98, 6.99, and 8.05 mm.

### 2.2 Statistical tests for outliers and group differences

Under the Lambertian assumption, radiance should not vary with distance or aperture. To evaluate this, statistical analyses were performed to identify outliers and group mean differences across each condition. Outlier detection was performed using Grubbs’ test (one-sided, α=0.05) and a Median Absolute Deviation (MAD) rule of thumb (using *k*=3 as the cutoff). For Grubbs’ test, the test statistic was compared to the critical value *G*_*critical*_, for *n* datapoints at α=0.05. A data point was flagged as an outlier if *G > G*_*critical*_. Separately, the MAD approach computed each point’s absolute deviation from the median. Any point whose deviation exceeded *k* times the robust deviation estimate was labeled as an outlier. To evaluate whether mean radiances differed significantly across distance or aperture settings, we performed a one-way ANOVA at α = 0.05. Each distance or aperture condition formed one “group,” with repeated measurements (triplicates) providing within-group variance. The resulting *F*-ratio and associated *p*-value were used to test the null hypothesis that all group means were equal.

### 2.3 Conversion of radiance into camera response

This section recreates the radiometric conversion formalism of Litorja^29^ to validate that the RET can serve as the calibrated source for SI-traceable radiance transfer under standard imaging geometries. Using the RET’s calibrated radiance output for conversion of the camera response involves imaging the RET emission aperture while accounting for the lens aperture size and the source-to-aperture distance. The relevant parameters and simplified 2D geometric schematics involved in the imager response conversion are shown in **Fig 3(a,b)**. The following derivations and equations are adapted from Litorja.^29^ Two optical geometries are considered: (1) imaging of the RET emission area onto the image sensor [**Fig 3(a)**] and (2) the aperture-limited light rays for the projection of a single source emission point onto the sensor [**Fig 3(b)**]. These two configurations define four solid angles determined by the source area (*A*_*src*_), image area (*A*_*img*_), aperture size (*A*_*apt*_), source-to-aperture distance (*d*) and aperture-to-image sensor distance (*d*_*sen*_). In the first geometry [**Fig 3a**] the two defined solid angles are 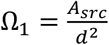 and 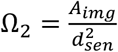 where Ω_1_ = Ω_2_. For the second optical geometry [**Fig 3b**] the two defined solid angles are 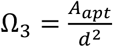 and 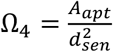. Note that the “source-to-aperture distance” *d* is measured from the emitting surface of the RET to the lens’s entrance pupil – the effective aperture position that defines the collection solid angle (Ω_3_) – rather than to the mechanical diaphragm location.

**Fig. 3:**
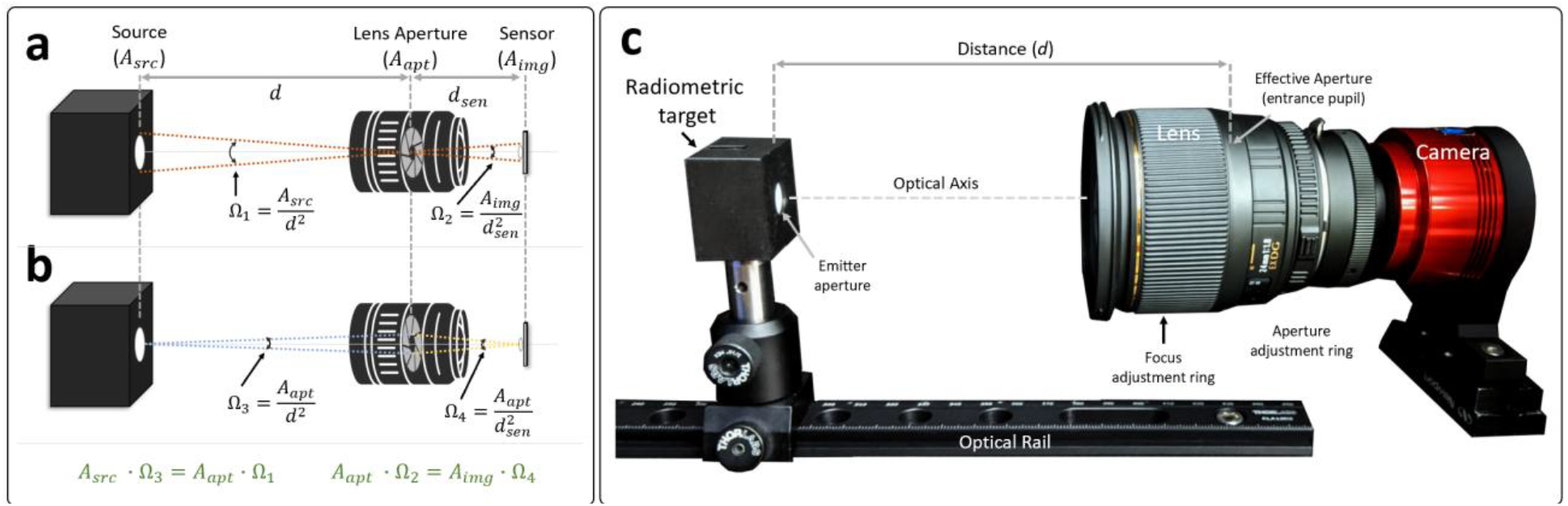
Optical set-up for assessing the radiance conversion of the camera response: **(a)** Geometry associated with imaging the RET source area onto the image sensor. **(b)** Geometry associated with the aperture-limited light rays for the projection of a single source emission point onto the sensor. **(c)** Experimental setup used to test the imager response conversion for varying distances and apertures.

As described in Litorja,^29^ the product of the area and solid angle *A* · Ω, often referred to as the *étendue*,^34^ remains constant in an optical system. This principle enables calculation of the radiant flux (Φ) passing from the RET through the lens aperture and projected onto the imaging sensor. The radiant flux, which is reduced by the lens transmittance, is projected onto the imaging sensor, such that:

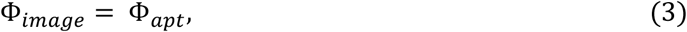

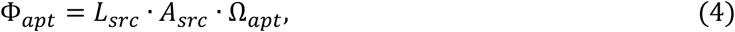

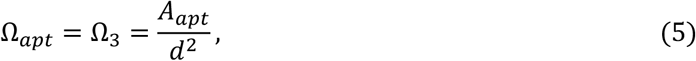

So that

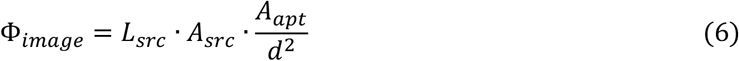

Hence, the measured radiant flux at the image plane *Φ*_*image*_ can be determined from the RET radiance (*L*_*src*_), RET source area (*A*_*src*_), lens aperture area (*A*_*apt*_), and source-to-aperture distance (*d*). From an imaging perspective, photons emitted by the RET are converted to electrons within each pixel, then digitized to a corresponding digital number *N* at each pixel (bound by the camera bit depth). The sum of the digitized outputs (*N*_*i*_) for all pixels (*i*) corresponding to the RET emitting area is noted as *S*, such that

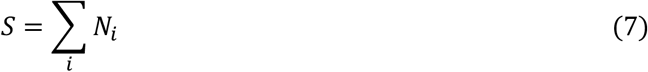

Lastly, the imager responsivity in the specified spectral band, *R*_*λ*_, is the ratio of total counts to the image-plane radiant flux:

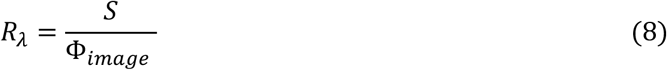

with units of counts/W. Hence, if the source is Lambertian and the principle of *étendue* holds, *R*_*λ*_ should remain invariant with both distance and aperture size – thereby enabling direct conversion of SI-traceable radiance into the camera response.

#### 2.3.1 Imager Responsivity vs. Distance

**Fig 3(c)** shows the experimental setup used to test imager responsivity *R*_*λ*_ with varying source-to-aperture distances (*d*). A 12-bit back-illuminated CMOS camera (Suzhou ZWO Co, ASI462MM) and lenses were used to image the RET emission surface with the RET optically aligned on an adjustable optical rail. The RAW image data (12-bit native digitization) were exported in 16-bit TIFF format and used for subsequent analysis. A Navitar 16mm lens (Navitar, Machine Vision Lens 16mm f1.4) and Sigma 24 mm lens (SIGMA Corporation, #432306) were used to test the imaged radiance invariance with distance by using a fixed aperture area (*A*_*apt*_) and varying the source-to-aperture distance (*d*). The Navitar images were acquired with a fixed f-number aperture of 4.0, gain of 80, exposure time of 10 ms, and *d* = 100, 150, 200, 300, 350, and 400 mm; the Sigma lens images were acquired with a fixed f-number aperture of 5.6, gain of 80, exposure time of 10 ms and *d* = 100, 150, 200, 300, and 350 mm. The exposure times were selected to ensure similar signal-to-noise performance and keep the imaging sensor within its linearity range (i.e., ∼80% of total well capacity). Images were acquired with a RET radiance output of 54 µW·cm^-2^·sr^-1^. Aperture size was determined by each lenses’ f-number *n* setting, defined as *n* = *f*/*D*, where *f* is the focal length of the lens and *D* is the aperture diameter. The corresponding aperture area *A*_*apt*_ can be calculated from:

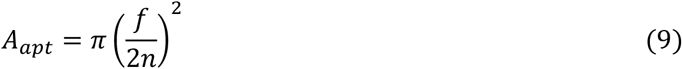

Therefore, the calculated aperture area *A*_*apt*_ for the Sigma and Navitar lens acquisitions were 14.4 mm^2^ and 12.6 mm^2^, respectively. Each lens was refocused at every tested *d* to ensure the RET emission surface remained in focus, thereby preserving image quality and measurement consistency across distances.

#### 2.3.2 Imager Responsivity vs. Aperture Size

To examine the invariance of the imager responsivity *R*_*λ*_ with respect to aperture settings, the aperture size (*A*_*apt*_) was varied at a fixed source-to-aperture distance (*d*). The Sigma 24mm lens and a Nikon 50mm lens (Nikon, #1433) were used. Both image sets were acquired with a RET radiance output of 54 µW·cm^-2^·sr^-1^ and f-number apertures settings of 2.8, 4, 5.6, 8, and 11. A fixed distance *d* of 300 mm and 400 mm were used for the Sigma and Nikon lenses, respectively. Imaging parameters were set to an exposure time of 9 ms and gain of 80, ensuring that the largest aperture setting (f/2.8) produced ∼80% sensor saturation to preserve signal linearity. Each corresponding aperture area was calculated using **Equation (9)**.

#### 2.3.3 Image and Responsivity Analysis

All images (12-bit native digitization) were acquired as uncompressed 16-bit TIFFs and analyzed in ImageJ to extract the total pixel counts (*S*) over the 10 mm RET emission surface. Imaged radiant flux values Φ_*image*_ were calculated using **Equation (6)**. The resulting *R*_*λ*_ was calculated using **Equation (8)**. To evaluate whether the mean imager responsivity differed significantly across distance or aperture settings, we performed a one-way ANOVA at α = 0.05. Each distance or aperture condition formed one “group,” with repeated measurements (triplicates) providing within-group variance. The resulting *F*-ratio and associated *p*-value were used to test the null hypothesis that all group means were equal.

### 2.4 Physical vs. Digital apertures for ROí radiance transfer

To extend the concept of imager responsivity (*R*_*λ*_) to digitally defined region-of-interest (ROI) and pixel-level analyses, we evaluated the *R*_*λ*_ for both physical apertures and “digital apertures.” For this, the Sigma and Nikon lenses were used to image the RET under three physical aperture conditions: (1) No aperture (RET emission surface of 10 mm ø), a 5 mm physical aperture (Thorlabs, P5000K), and 2 mm physical aperture (Thorlabs, P2000K). All images were acquired using a RET radiance of 54 µW·cm^-2^·sr^-1^, camera gain of 80, an integration time of 30 ms, and an f-number of 5.6, with the Sigma lens positioned at 300 mm and the Nikon lens at 400 mm. The ZWO camera was used for imaging. By applying digital masks (ROIs) to these images in post-processing, the variability of *R*_*λ*_ with both physical aperture size and smaller digitally defined regions was assessed. All images (12-bit native digitization) were acquired as uncompressed 16-bit TIFFs and analyzed in ImageJ.

### 2.5 Radiance Transfer Curve (RTC) for imager characterization

A key limitation of relying solely on the imager responsivity *R*_*λ*_ is that it assumes a strictly linear relationship between imaged radiance and pixel intensity. In practice, imaging sensors exhibit non-linear behavior, particularly at the low and high signal regimes. Additionally, image-processing pipelines – including bit-depth compression, logarithmic or gamma scaling, baselining, among others – can introduce further non-linearities. As a result, many fluorescence imaging systems, including those used in FGS, may produce outputs that do not scale linearly with radiance and cannot be readily compared across systems.

To address these complexities, we borrow principles from imaging photon transfer^40^ and introduce the *Radiance Transfer Curve* (RTC) to incorporate sensor and image processing non-linearities into the conversion of imager response into SI-traceable units. The acquisition of the RTC involves imaging a Lambertian source at various SI-traceable radiances (*L*_*cal*_) while holding the critical imaging parameters – exposure time, gain, aperture size *A*_*apt*_, aperture-to-image-sensor distance *d*_*sen*_, optical components, and image processing pipeline – fixed [**Fig 4(a)**]. These radiances span both the noise floor and saturation point (i.e full-well capacity) of the imaging sensor. The resulting image set [**Fig 4(b)**] is analyzed to extract a mean pixel intensity value over the Lambertian source’s emission region, generating a look-up-table (LUT) which we term the RTC. Plotting the mean intensities versus calibrated RET radiance output [**Fig 4(c)**] visualizes the RTC for the specified spectral band and imaging configuration.

**Fig. 4:**
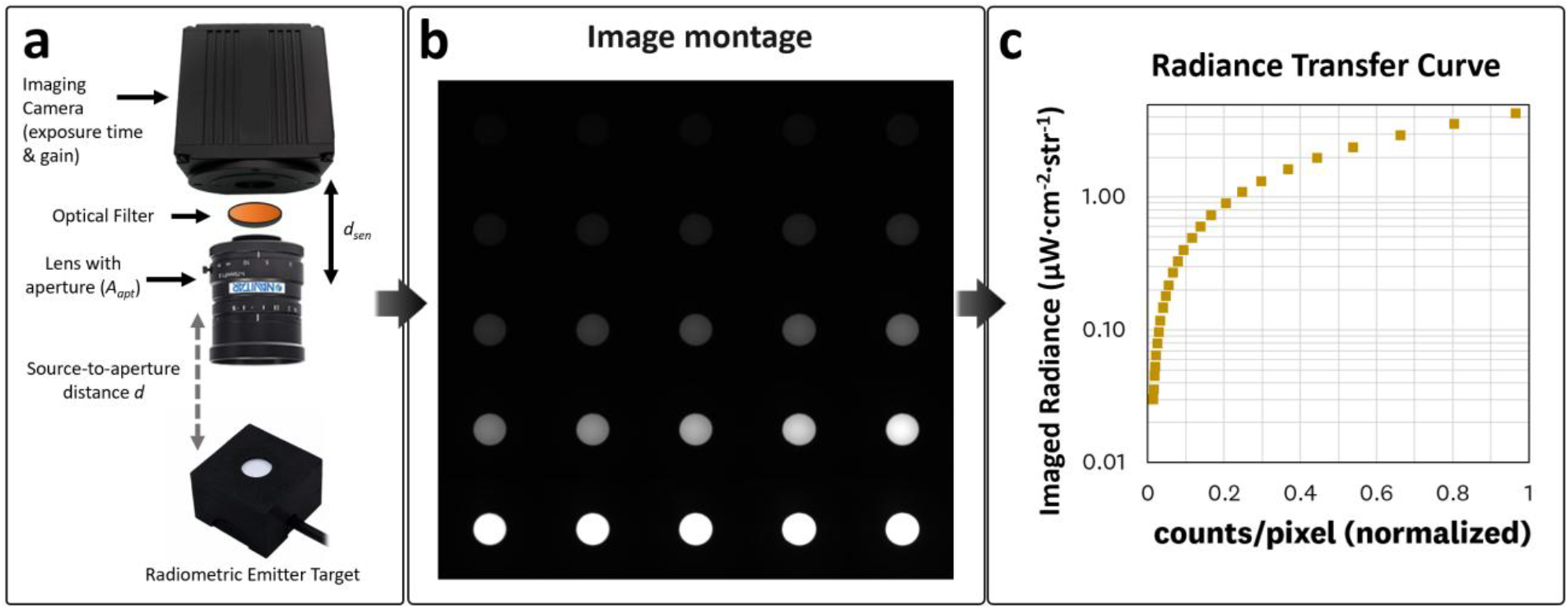
Acquisition of the RCT for a given imaging configuration: **(a)** Schematic showing a simplified imaging set-up for characterizing of the *Radiance Transfer Curve (*RTC) alongside the significant parameters **(b)** Example images acquired at different radiances, from which mean pixel intensities are computed. **(c)** Plot of imaged radiance vs. mean pixel intensity, yielding the RTC.

A key insight – mathematically formalized in Appendix A – is that for a Lambertian source imaged through a lens system, the mean pixel intensity, *I*_*mean*_, is independent of the source-to-aperture distance *d*. As summarized by the relationship

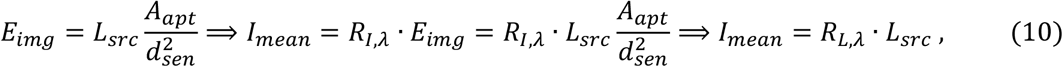

where the imaged flux is distributed over a correspondingly smaller or larger area on the detector such that the imaged irradiance *E*_*img*_ is related only to the lens aperture area *A*_*apt*_, lens-detector distance *d*_*sen*_, and source radiance *L*_*src*_. Consequently, the RTC measurement can be performed at any distance *d*, provided the Lambertian source is sufficiently resolved in the acquired image. Here, *R*_*I,λ*_ denotes the irradiance-to-counts responsivity factor for spectral band λ (units: counts·μW^-1^·cm^2^) and, given that *A*_*apt*_ and *d*_*sen*_ are normally fixed for a given imager configuration, we define the radiance-to-counts responsivity factor *R*_*L,λ*_= *R*_*I,λ*_·*A*_*apt*_·*d*_*sen*_^*-2*^ Thus, with *A*_*apt*_, *d*_*sen*_, exposure time, and gain held fixed, any measured pixel intensity can be converted directly to radiance across the image plane, without additional distance corrections. In other words, we can directly relate the calibrated radiance values *L*_*cal*_ to the mean pixel intensity *I*_*mean*_ given fixed imaging parameters (*A*_*apt*_, *d*_*sen*_, exposure time, gain) such that *I*_*mean*_ = *R*_*L,λ*_ · *L*_*Cal*_ where *R*_*L,λ*_ is the RTC look-up-table entry relating pixel intensity to radiance for a specific spectral band, with units of counts·µW^-1^·cm^2^·sr^1^. By sampling a range of *L*_*cal*_ values spanning the sensor’s dynamic range, we build the LUT of *R*_*L,λ*_ values. This RTC then provides an SI-traceable conversion from pixel intensities to the calibrated imaged radiance.

Because of the distance invariance, the same RTC can be applied over a broad range of working distances *d*, so long as the optical configuration (i.e., *A*_*apt*_, *d*_*sen*_, exposure time, and gain) remains unchanged. For a given system, multiple RTCs can be acquired to account for different system settings or configurations. Additionally, the RTC inherently captures aggregate effect of image-processing pipelines and optical elements, further extending its usability. Once acquired, the resulting RTC is used in Section 2.6 to perform the radiance and Radiance Imaging Transforms.

#### 2.5.1 Capturing the RTC and assessing R_λ_ nonlinearities

To evaluate the RTC acquisition, a 12-bit sCMOS camera (Tucsen, FL-20BW), 800 LP filter (Thorlabs, FELH0800), and the Sigma 24mm lens (with aperture set to *f/*5.6) were used to image the RET at 49 different radiances ranging from 0.04-225.7 µW·cm^-2^·sr^-1^. A gain of 350 and exposure time of 200 ms were used. Mean pixel values were extracted from the corresponding ROI of the 10 mm diameter RET emission region from 16-bit RAW (.TIFF) images using ImageJ. The calibrated radiance values *L*_*cal*,_ obtained as described in Section 2.1, are plotted against the mean pixel intensity *I*_*mean*_ resulting in the RTC curve for this imaging configuration.

### 2.6 Radiance Imaging Transform (RIT) and Fluorescence Imaging Transform (FIT)

Having acquired the RTC for a particular spectral band and imaging configuration (Section 2.5), we now describe its use to convert fluorescence signals from digital counts (*N*) to SI-traceable radiance (µW·cm^-2^·sr^-1^). We term this process the *Radiance Imaging Transform* (RIT). In brief, the RIT converts the digital counts of an image [**Fig 5(a)**] into imaged radiance using the RCT as a LUT for each pixel [**Fig 5(b)**], resulting in the RIT image [**Fig 5(c)**] with SI-traceable units of µW·cm^-2^·sr^-1^. This process transforms the digital counts of each pixel (e.g. 8-bit, 12-bit, 16-bit) into a float-32 number with corresponding radiance units. We next introduce the *Fluorescence Imaging Transform* (FIT), which extends this framework by performing a pixel-by-pixel normalization of the RIT image to the measured excitation irradiance, *E*_*exc*_*(x,y)*, where *(x,y)* denote spatial coordinates in the image. This normalization can account for spatial non-uniformities in excitation and converts the imaged radiance to aggregate fluorescence yield with units of sr^-1^,^29^ resulting in the FIT image [**Fig 5(d)**]. The following subsections explain the methods associated with the generation of RIT and FIT images and the evaluation of their performance using a fluorescence concentration reference target and a breast lumpectomy phantom.

**Fig. 5:**
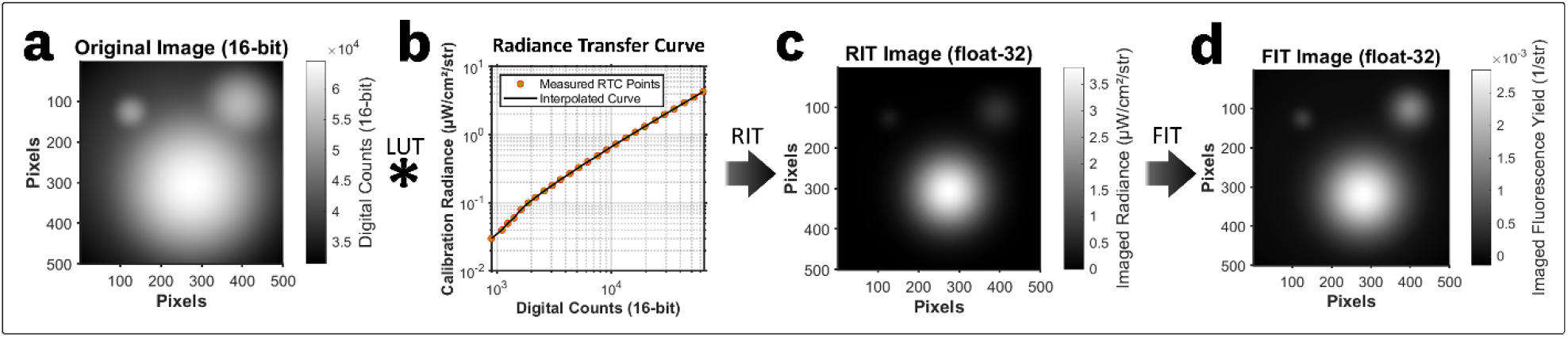
The Radiance Imaging Transform (RIT) and Fluorescence Imaging Transform (FIT) enable conversion of fluorescence images to SI-traceable radiometric units. (a) Simulated 16-bit fluorescence image (500 × 500 pixels) with a gamma curve containing three computationally generated Gaussian distributions, representing embedded fluorescent features captured under a defined optical configuration with a broad, spatially varying Gaussian illumination profile. The corresponding radiance transfer curve (RTC) of the imaging system, established via calibration with a Lambertian radiance source, serves as a pixel-wise look-up table (LUT) to map digital counts to imaged radiance. (c) Application of the RTC to each pixel yields the Radiance Imaging Transform (RIT) image (float-32) with units of radiance (µW·cm^-2^·sr^-1^). (d) Pixel-wise normalization of the RIT image by the excitation irradiance, *E*_*exc*_*(x,y)*, results in the aggregate fluorescence yield FIT image with corresponding units of sr^-1^.

#### 2.6.1 RIT at the pixel or ROI Level

The RIT converts each pixel’s digital counts, *I*_*pixel*_ (or mean pixel counts within an ROI), into an imaged radiance, *L*_*img*_ (µW·cm^-2^·sr^-1^), by applying the RTC as a radiance LUT. Because an RTC is established under fixed optical and camera settings (exposure time, gain, aperture size *A*_*apt*_, aperture-to-image-sensor distance *d*_*sen*_, etc.), it inherently accounts for sensor non-linearities, optical components, and image-processing pipelines. The relationship is given by:

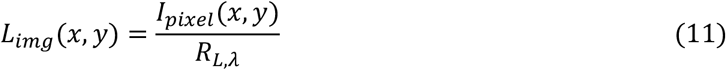

where *R*_*L,λ*_ is the LUT entry relating pixel intensity to radiance in the relevant spectral band. Here, *(x,y)* denote spatial coordinates in the detector (i.e., image) plane, corresponding to the horizontal and vertical pixel positions of the sensor array. Section 3.3 results show that imaging system response remains valid for smaller digital ROIs, ensuring that each pixel – or sub-ROI – can be treated independently for fluorescence radiance transfer. Moreover, given the distance invariance of the RTC, the RIT can be applied at arbitrary source-to-aperture distances (i.e., working distances) without additional corrections. Since the RTC is collected at discrete radiance values, interpolation is required to map each pixel or ROI (*I*_*pixel*_ or *I*_*mean*_, respectively) to corresponding imaged radiances (*L*_*img*_) [**Fig 5(b)**]. The resulting RIT image [**Fig 5(c)**] contains pixels of float-32 data with units of µW·cm^-2^·sr^-1^. Subsequently, this RIT image can be normalized by the excitation irradiance at the imaging plane to produce the FIT image [**Fig 5(d)**], where each pixel of float-32 data is mapped to an aggregate fluorescence yield with corresponding units of sr^-1^. The FIT is further discussed in Section 2.6.2. It is worth noting that nonlinear sensor response and image-processing transformations can alter ROI-averaged values; therefore, applying the RIT at the pixel level is preferred over ROI-level conversion to avoid systematic bias and minimize artifacts.

#### 2.6.2 FIT Excitation Normalization

Although the RTC-based RIT is distance-invariant for Lambertian emitters (Section 2.5, Appendix A), the excitation irradiance incident on the fluorescence imaging plane typically varies with working distance, illumination profile, excitation beam divergence, and other optical parameters. To account for these excitation-dependent variations, the imaged radiance *L*_*img*_ can be normalized by the measured excitation irradiance *I*_*exc*_ at the imaging plane, resulting in a quantity termed aggregate fluorescence yield. We extend this concept to the RIT methodology and introduce the FIT image which has corresponding units of sr^-1^.

The FIT is an expansion of the aggregate fluorescence yield factor *F*, previously defined by Litorja,^29^ where the imaged fluorescence emission radiance *L(λ*_*em*_) is normalized by the excitation irradiance *E(λ*_*ex*_), such that

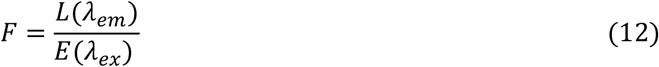

Unlike the molar fluorescence yield, which represents the fluorescence efficiency of a pure fluorophore, the aggregate fluorescence yield *F* incorporates additional modifying factors such as fluorophore concentration, absorbance at the excitation wavelength, and surrounding optical properties. Specifically, the aggregate fluorescence yield represents the effective fluorescence efficiency of the sample in response to a known excitation irradiance.

We extend this concept by defining the FIT as the per-pixel aggregate fluorescence yield at emission wavelength (or band) *λ*_*em*_ under excitation *λ*_*ex*_, expressed as

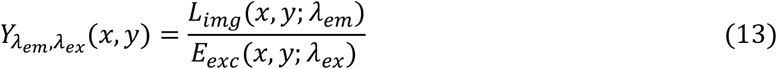

which implements the FIT as a pixel-wise normalization of the RIT radiance image *L*_*img*_ by the corresponding excitation *E*_*exc*_ measured at the imaged object plane. Here, (x,y) denote spatial coordinates in the detector plane, corresponding to the horizontal and vertical pixel positions of the sensor array. For systems exhibiting approximately homogeneous excitation across the field of view, a scalar (field-averaged) normalization may be used in practice, such that

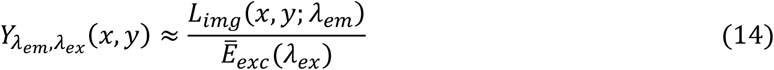

Theoretically, FIT enables direct cross-system comparisons by minimizing the impact of instrument-specific response and distance-dependent fluorescence excitation effects. In other words, a properly acquired FIT image should map the acquired fluorescence signal into a shared, absolute fluorescence space that is independent of specific imaging parameters, excitation irradiance, and image-processing pipelines. Consequently, FIT images acquired from different fluorescence imaging systems or varied illumination geometries should become directly comparable, facilitating cross-system validation and the development of standardized quantitative fluorescence imaging.

#### 2.6.2 Assessing the RIT for fluorescence over a wide dynamic range

To assess the use of the RIT for specific fluorescence bands and over a wide dynamic range of fluorescence intensities we utilized an ICG-equivalent concentration sensitivity target (RCS-ICG-ST01-QUEL02, QUEL Imaging). In brief, the RCS target uses tissue-equivalent optical formulations (µ_a_ = 0.021 mm^-1^, µ_s_’ = 0.27 mm^-1^ at 800 nm) in 9 wells with varying ICG-equivalent concentrations (1-1000 nM alongside a control well) and serves as a shelf stable fluorescence signal sensitivity target over a wide range of fluorescence signal levels. The fluorescence emission output of this phantom closely mimics that of ICG in plasma^18,22^ such that the RET emission spectrum [**Fig 1(c)**] provides good spectral responsivity overlap. This overlap allows the RIT methodology, using the acquired RTC from Section 2.5.1, to convert the acquired RAW image into imaged ICG emission radiance (µW·cm^−2^·sr^−1^). The acquired 12-bit RAW fluorescence image (785 nm laser excitation, 800 nm long-pass emission collection) of the concentration sensitivity target was analyzed by using 5 mm ROIs centered on each 10 mm well to create a plot of the mean pixel intensity *I*_*mean*_ vs. ICG concentration. The 12-bit image was converted into units of imaged radiance *L*_*img*_ using the RIT (Section 2.6.1) with identical 5 mm ROIs centered on each 10 mm well utilized to generate the *L*_*img*_ vs. ICG concentration. The RIT image conversion was performed in MATLAB using the *interp1* function with shape-preserving piecewise cubic interpolation. The fluorescence imaging configuration, including the RCS, RET, and irradiance power meter, are shown in **Fig 6**, with imaging system component details provided in Section 2.5.1.

**Fig. 6:**
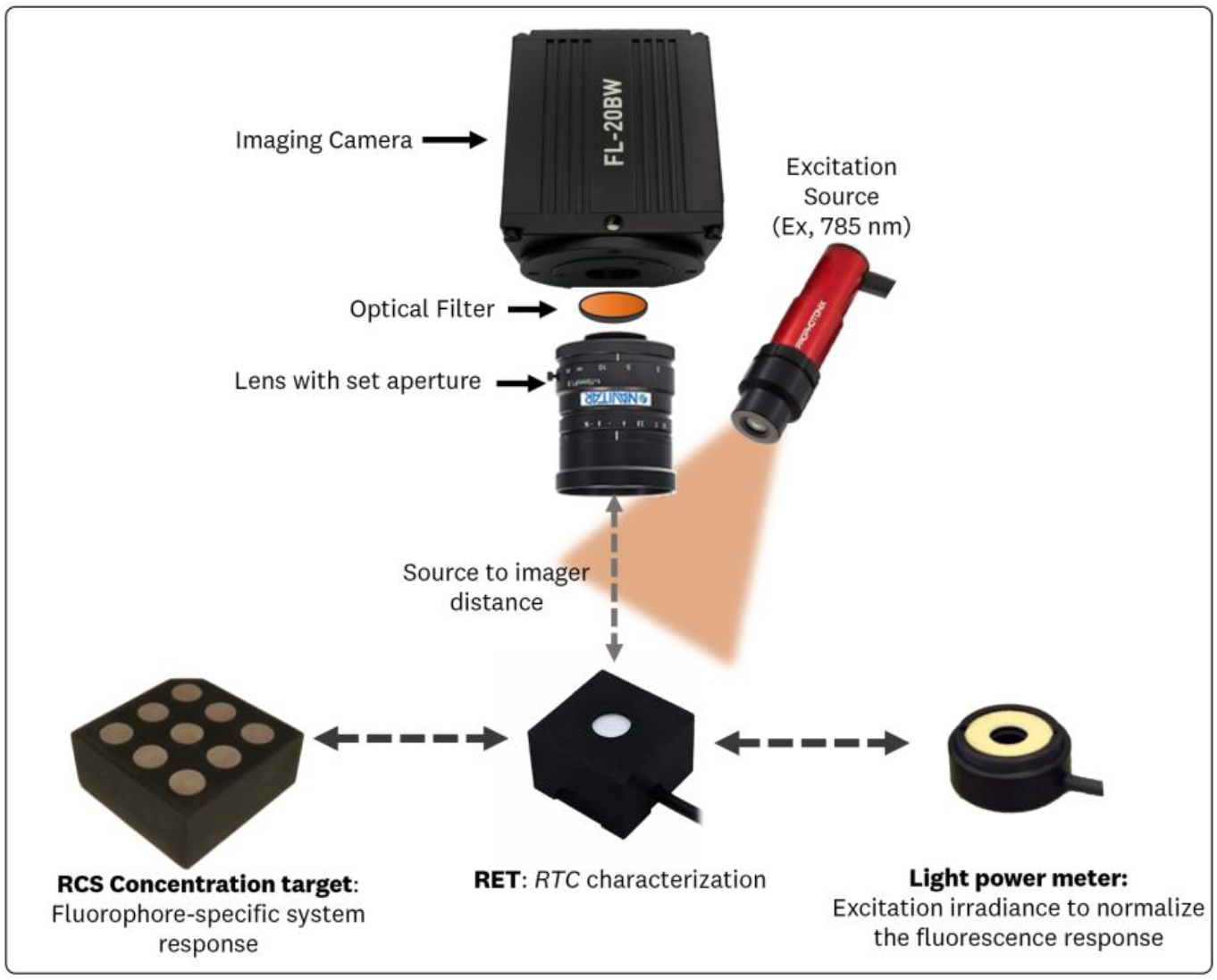
Fluorescence imaging configuration for measuring the reference concentration sensitivity target, radiance transfer curve (RTC) of the optical configuration, and the excitation irradiance

#### 2.6.3 Assessing the RIT using varying image processing pipelines

To assess the effectiveness of the RIT to account for image processing pipelines we transformed the RAW 12-bit images of the concentration sensitivity target (Section 2.6.2) and the RTC characterization (Section 2.5.1) through two image processing pipelines: 1) an 8-bit down-sampled image with baselining and 2) a log10 compression that is re-mapped into a 12-bit image space. These pipelines mimic traditional down-sampling that is used for image display and generation in FGS devices^16,18,19^ and the extension of the visualized dynamic range to better align with human brightness perception,^48^ respectively. The *I*_*mean*_ of 5 mm ROIs centered on each well of the RCS target for the three pipelines (RAW, 8-bit, and log10) were extracted from the digital count images. Subsequently, the RTC for each respective image processing pipeline was used to perform the RIT on each image to obtain the mean *L*_*img*_ of the 5 mm ROIs centered on each well. The RIT image conversion was performed in MATLAB using the *interp1* function with shape-preserving piecewise cubic interpolation. To better understand the use of the RIT to converge fluorescence reporting metrics across varying image processing pipelines, four metrics were calculated for each fluorescent well of the concentration target using the mean and standard deviation identified by each of these methods: (1) signal-to-background ratio (SBR), (2) signal-to-noise ratio (SNR), (3) contrast-to-noise ratio (CNR), and (4) contrast-to-variability ratio (CVR).^49,50^ The metrics are calculated as 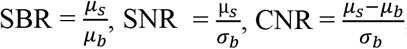, and 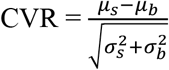, where *μ*_*b*_ and *σ*_*b*_ are the mean and standard deviation of the signal ROI (1000-1 nM wells), and *μ*_*b*_ and *σ*_*b*_ are the mean and standard deviation of the background ROI (control well).

#### 2.6.4 Breast Lumpectomy Phantom Imaging

The RIT was further assessed on a breast lumpectomy phantom, imaged with the system described in Section 2.5.1, to test its applicability in anthropomorphic geometries with tumor inclusions at ICG-equivalent concentrations of 30, 100, 300, and 1000 nM. The breast lumpectomy phantom fabrication methods have been described previously, with measured absorption and reduced scattering coefficients of 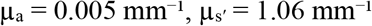, and 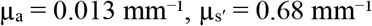 at 800 nm, for the bulk and tumor material, respectively.^50,51^ The bulk breast material was manufactured with 3 nM ICG to mimic background tissue autofluorescence. The RTC acquired in Section 2.5.1 was used to perform the RIT, converting the acquired 12-bit RAW images to imaged ICG emission radiance (µW·cm^−2^·sr^−1^). The RIT image conversion was performed in MATLAB using the *interp1* function with shape-preserving piecewise cubic interpolation. The excitation irradiance was measured, using the S120VC photodiode, at the imaging plane and used to generate the FIT image (sr^−1^). The field-averaged **Equation (14)** was used given the approximately homogeneous excitation across the phantom field of view. Contrast metrics of SBR, SNR, CNR, and CVR, as defined in Section 2.6.3, were computed from 5 mm ROIs, as shown in **Fig 14(b)**.

## 3 Results

### 3.1 Lambertian emission of the radiometric target

The results of evaluating the RET’s Lambertian emission are shown in **Fig 7**, which compares measured radiance under two conditions: (a) varying source-to-aperture distance and (b) varying aperture size. Statistical analyses included Grubbs’ and Median Absolute Deviation (MAD) outlier tests, as well as one-way ANOVA across each set of mean radiance values. The methods of these measurements and analysis are covered in Section 2.2.

**Fig. 7:**
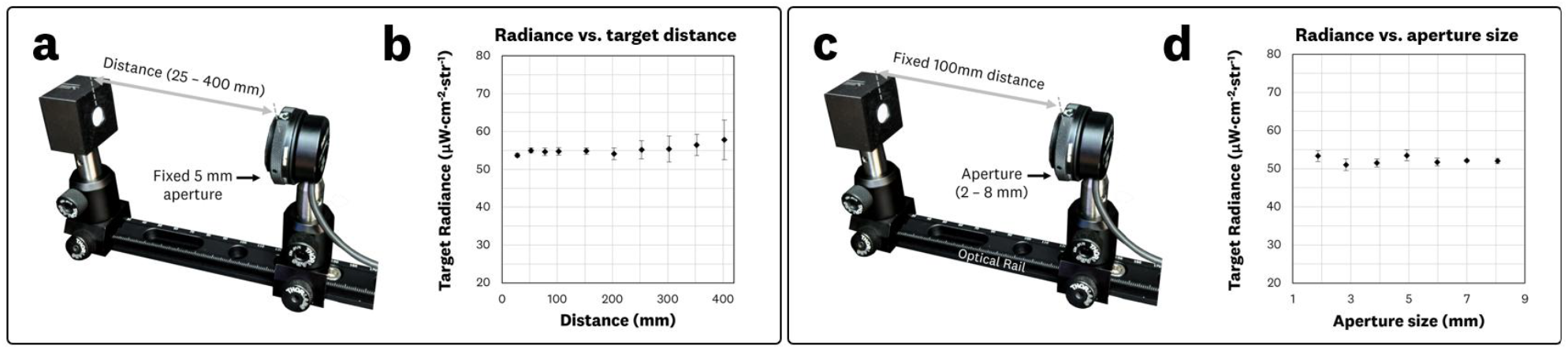
Results of the Lambertian emission evaluation of the RET: **(a)** varying distance experimental set-up and (b) plot of the measured radiance vs. distance data. **(c)** Varying aperture experimental set-up and **(d)** plot of the measured radiance vs. aperture size data.

In the distance-varying experiment [**Fig 7(a,b)**], no outlier was flagged by Grubbs’ test (*n=*10, *G*_*max*_ = 2.28 < *G*_*critical*_ = 2.82). However, the largest distance (400 mm) was identified as an outlier by the MAD criterion (400 mm radiance = 57.7 > cut-off value = 56.7 µW·cm^-2^·sr^-1^), likely due to the signal approaching the detector’s lower detection limit at this distance. Additionally, a trend of increasing standard deviation was observed as distance increased, corresponding to a decline in signal-to-noise ratio; for instance, the 400 mm measurement had approximately 200× less signal than the 25 mm measurement. Despite this discrepancy, a one-way ANOVA across all tested distances showed no significant difference (p ≈ 0.88), indicating radiance remained effectively constant with distance.

For the aperture-varying experiment [**Fig 7(c.d)**], neither Grubbs’ test (*n=*7, *G*_*max*_ = 1.43 < *G*_*critical*_ = 2.02) nor MAD criteria (max deviation = 1.47 < MAD cut-off = 2.38 µW·cm^-2^·sr^-1^) flagged any outlier. A one-way ANOVA also did not reveal any statistically significant difference (*p* = 0.15) among aperture settings. These findings confirm that the measured radiance remained effectively constant with the solid angle subtended by the aperture.

Overall, the combined outlier and ANOVA results demonstrate that the RET emission remains consistent over the tested distances and apertures. This Lambertian behavior is crucial for converting imager responses into SI-traceable radiances providing the necessary radiance invariance to perform imager response conversion into SI-traceable units.

### 3.2 Conversion of radiance to camera response

Having established the Lambertian emission of the RET (Section 3.1), we next evaluated the conversion of radiance into imager response according to the methods in Sections 2.3, 2.3.1, and 2.3.2. Specifically, we computed the imager responsivity, *R*_*λ*_, using **Equations (6)-(8)** for different source-to-aperture distances (*d*) and for varying aperture sizes (*A*_*a*pt_). **Fig 8** and **Fig 9** summarize these results.

**Fig. 8:**
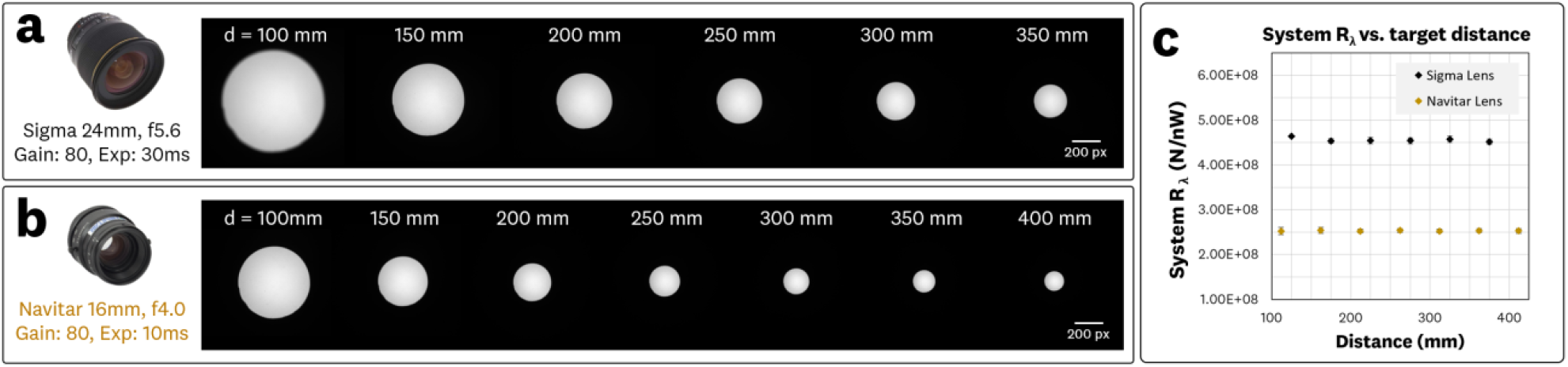
Results of the imager responsivity *R*_*λ*_ vs. source-to-aperture distance *d* measurements: **(a)** Sigma lens varying distance acquired images, **(b)** Navitar lens varying distance acquired images, and (c) the plot of the resulting system responsivity *R*_*λ*_ vs. source-to-aperture distance *d* measurements. All images share the same look-up-table scaling to provide cross-image comparison.

**Fig. 9:**
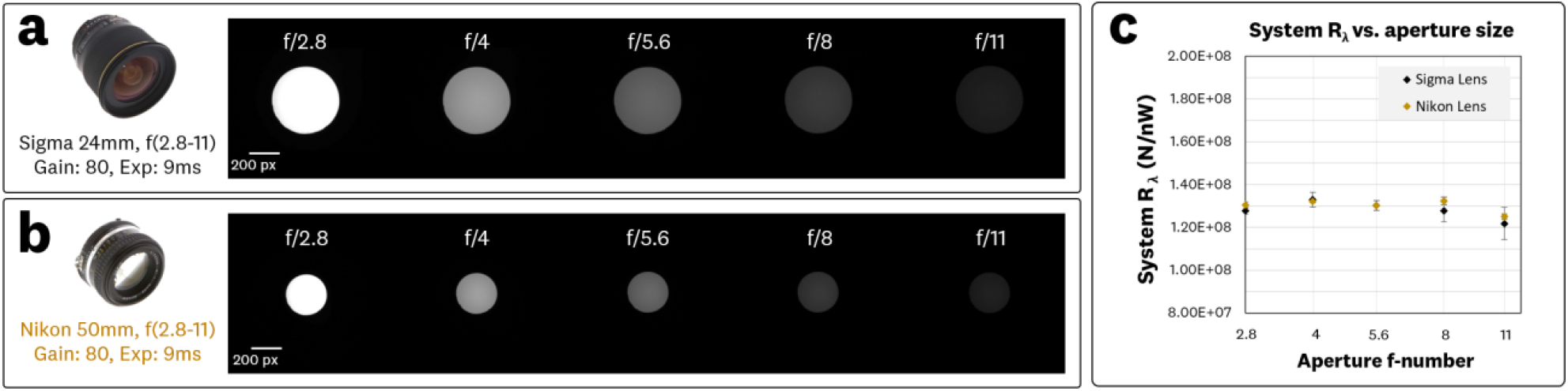
Results of the imager responsivity *R*_*λ*_ vs. aperture size comparison: **(a)** Sigma lens varying aperture acquired images, **(b)** Nikon lens varying aperture acquired images, and **(c)** the plot of the resulting system responsivity *R*_*λ*_ vs. aperture f-number plots measurements. All images share the same look-up-table scaling to provide cross-image comparison

#### 3.3.2 Imager responsivity vs. distance

**Fig 8** shows the acquired images for the Sigma 24 mm lens [**Fig 8(a)**] and Navitar 16 mm lens [**Fig 8(b)**] alongside the plotted *R*_*λ*_ vs *d* data [**Fig 8(c)**]. All images were displayed using a common look-up-table scaling (0–16-bit range mapped to black and white, respectively) to enable direct cross-image comparison. Both lenses were used to image the RET at distances *d* of 100–400 mm with fixed apertures. The measured *R*_*λ*_ remained consistent across all tested distances, with average values of approximately 4.47 counts/W [**Fig. 8(a)**] and 2.48 counts/W [**Fig. 8(b)**] for the Sigma and Navitar lens acquisitions, respectively. A one-way ANOVA indicated no statistically significant difference among these mean values (p = 0.48 for the Sigma lens; *p* > 0.99 for the Navitar lens). These findings confirm that, under the tested conditions, the imager response scales appropriately with the RET radiance regardless of distance, in line with the Lambertian and *étendue* assumptions.

The difference in the absolute *R*_*λ*_ between the two lenses reflects variations in imaging parameters (aperture setting, exposure time) as well as differences in lens transmission and optical design. Nonetheless, each system’s *R*_*λ*_ remained stable over the examined distance range, supporting the expected invariance of responsivity with distance for a Lambertian source.

#### 3.2.2 Imager responsivity vs. aperture size

**Fig 9** shows the acquired images for the Sigma [**Fig 9(a)**] and Nikon [**Fig 9(b)**] lenses alongside the plotted *R*_*λ*_ vs aperture size data [**Fig 9(c)**]. All images share the same look-up-table scaling to provide cross-image comparison; the scaling used was normalized to the imaging set maxima. Identical camera settings were utilized with fixed distances and varied apertures as described in Section 2.3.2.

For the Sigma Lens dataset [**Fig 9(a)**], a one-way ANOVA provided *p* = 0.10 when all five apertures measurements were included, indicating no statistically significant difference in responsivity. Excluding the f/11 measurement (where small manufacturing tolerances can have a larger relative impact) increased *p* to 0.26. For the Nikon Lens dataset [**Fig 9(b)**], the ANOVA provided *p* = 0.001 when all f-numbers were included, but rose to 0.45 when the f/11 datapoint was removed. This improvement suggests that the largest f-number measurement can be sensitive to minor lens-aperture deviations (e.g., manufacturing tolerances of ±100 µm) that become more proportionally significant at smaller aperture diameters; these manufacturing deviations, along with the inability to measure the true lens aperture size, can alter the statistical outcome of the ANOVA test. Similar observations have been reported previously by Litorja.^29^

Overall, these aperture-variation results are consistent with the theoretical expectation that, under Lambertian conditions and *étendue* assumptions, *R*_*λ*_ should not depend on the lens aperture. Where minor deviations did arise (aperture settings of f/11), they appear attributable to practical manufacturing tolerances rather than a violation of the *étendue* principle.

### 3.3 Physical vs. Digital apertures for radiance transfer

Having confirmed the Lambertian behavior of the RET (Section 3.1) and validated the conversion of radiance to imager response (Section 3.2), we next examined whether digital region-of-interest (ROI) selection could reliably replace physical apertures when determining the imager responsivity *R*_*λ*_. This approach is relevant to workflows where ROI-based measurements are used to isolate a sub-area of the imaged field. Unless otherwise noted, all *R*_*λ*_ values in this section are expressed in units of x10^8^ counts/nW.

**Fig 10(a)** shows representative images of the RET acquired with no physical aperture (10 mm emission area), a 5 mm physical aperture, and equivalent digital ROI. **Fig 10(b)** and **Fig 10(c)** present the resulting *R*_*λ*_ values for the Sigma and Nikon lens imaging sets, respectively. For the Sigma lens set [**Fig 10(b)**], the measured *R*_*λ*_ for the physical vs. digital 5 mm aperture was (4.8 ± 0.1) vs. (4.8 ± 0.1), and for the 2 mm aperture (5.0 ± 0.3) vs. (5.0 ± 0.1). Similarly the Nikon lens set [**Fig 10(c)**] yielded (5.1 ± 0.1) vs. (5.1 ± 0.1) for the 5 mm aperture and (5.2 ± 0.2) vs. (5.3 ± 0.1) for the 2 mm aperture. Given the difference in mean responsivity values for the 2 mm Nikon data, a two-tailed Welch t-test was performed (independent samples; N = 1,800 pixels; assumed spatial correlation factor = 0.05), which yielded *p* = 0.32, indicating no statistically significant difference in responsivity. These results indicate that there was no statistical difference between physically apertured measurements and those obtained via digital masking, which signifies that adjacent illuminated regions did not contribute substantially to the ROI’s measured radiance.

**Fig. 10:**
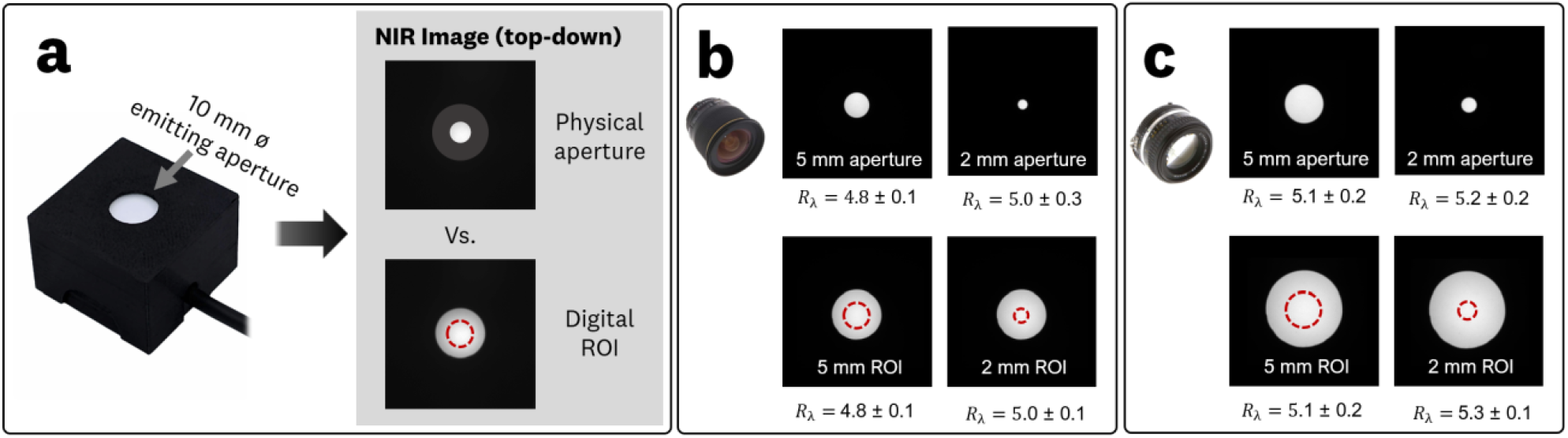
**(a)** Physical vs. digital ROI system responsivity experiment utilizing 5 mm and 2 mm apertures with respective images and calculated *R*_*λ*_ values for the imaging configurations utilizing the **(b)** Sigma 24 mm lens, and **(c)** Nikon 50 mm lens.

For the full 10 mm emission region, the measured *R*_*λ*_ values were 4.4 ± 0.3 (Sigma image) and 4.7 ± 0.4 (Nikon image). These values are slightly lower than those for the smaller apertures, reflecting the pseudo top-hat emission profile of the RET [**Fig 1(e)**]: the smaller central regions tend to have a more uniform, slightly higher radiance than the periphery. Although this difference does not alter the conclusion that physical and digital apertures provide equivalent results, it highlights the need to account for potential non-uniformities in the calibrated solid-state emitter when deriving *R*_*λ*_. This indicates that, in practice, responsivity values tied to the RET’s calibrated radiance *L*_*cal*_ should be derived from total digital counts *S* over the full 10 mm ROI to account for the pseudo-top-hat profile and minimize bias from peripheral roll-off [**Equations (7)**,**(8)**].

### 3.4 Radiance Transfer Curve (RTC) for imager characterization

The RET was imaged at 49 different radiances ranging from 0.3 to 225.7 µW·cm^-2^·sr^-1^ with the results shown in Figure 10. At *L*_*cal*_ ∼ 155 µW·cm^-2^·sr^-1^, which corresponded to ∼50% of the camera’s full-well capacity, an imager responsivity *R*_*λ*_ = 2.51x10^9^ counts/nW was measured. To explore the non-linearities associated with imager responsivity extrapolation, this *R*_*λ*_ value was used calculate the imaged radiant flux *Φ*_*image*_ transfer [**Equation (6)**] of the image set given the 24 mm lens, f/5.6 aperture, and 270 mm source-to-aperture. **Fig 10(b)** shows the radiant flux vs. voltage plot for the RET alongside the camera detected radiant flux *Φ*_*image*_ utilizing *R*_*λ*_ = 2.51x10^9^ counts/nW. Although the data follows the expected saturation at full-well capacity, a log-linear view [**Fig 11(c)**] further reveals significant deviation from the predicted linear behavior in the lower signal regime with ∼2x difference between the calculated and actual radiant flux value at the 1.1V setting.

**Fig. 11:**
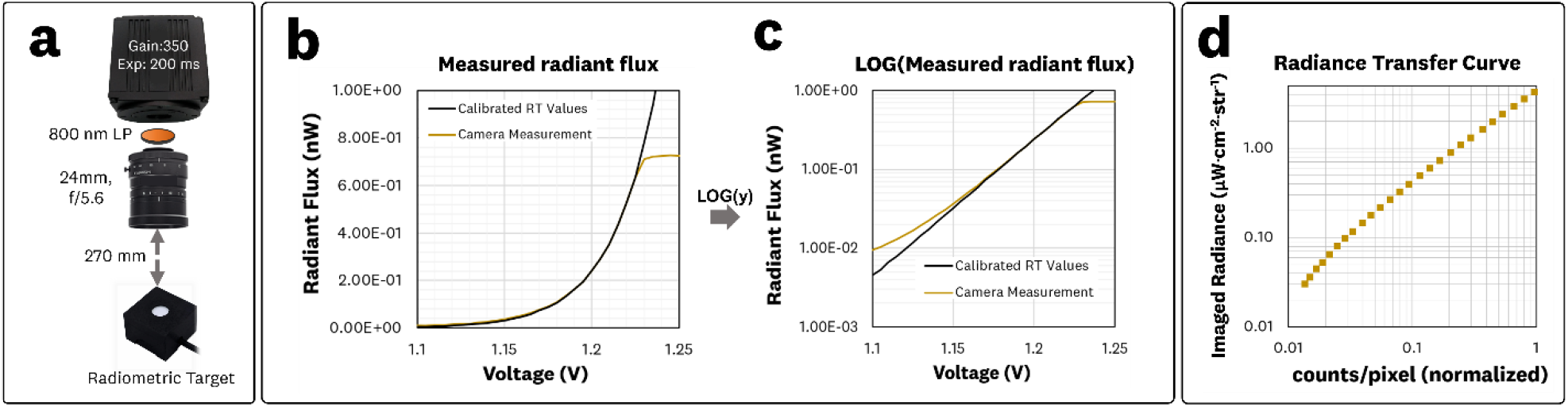
**(a)** Schematic of the imaging set-up for acquiring images for varying RET radiances in the 0.3 to 225.7 µW·cm^-2^·sr^-1^ range. **(b)** RET radiant flux values and camera-derived flux using the measured *R*_*λ*_ = 2.51x10^9^ counts/nW. **(c)** Log-linear plot of the radiant flux data showing significant deviation from linearity for the lower signal regime. **(d)** The resulting RTC plotting *L*_*cal*_ vs. mean pixel intensity *I*_*mean*_ (normalized) across the sensor’s dynamic range.

These results motivated the more comprehensive RTC approach which does not rely on a single *R*_*λ*._ to characterize the system response. Instead, the RTC samples a broad range of radiance values and directly correlates the mean pixel intensity *I*_*mean*_ with the known *L*_*cal*_ radiance of the RET. As described in **Section 2.5** and derived in **Appendix A**, we can relate *L*_*cal*_ = *I*_*mean*_/*R*_*L,λ*_ where *R*_*L,λ*_ is determined over multiple values of *L*_*cal*_, capturing both low and high-signal behavior. **Fig 11(c)** shows the resulting RTC for the given image configuration over the system’s dynamic range. This acquired RTC is used in **Section 3.5** and **Section 3.6** to produce the RIT and FIT transforms for imaging the ICG concentration target and the breast lumpectomy phantom.

### 3.5 Fluorescence Intensity Transform (RIT) and Fluorescence Imaging Transform (FIT)

Furthermore, an ICG concentration sensitivity target^22,51^ was imaged with the optical configuration detailed in Section 2.5.1. The collected fluorescence image of the concentration target is shown in **Fig 12(a)**. An excitation irradiance of 1.5 mW/cm^2^ was measured at the imaging plane. The corresponding average ROI intensity *I*_*mean*_ vs. ICG concentration plot is provided in **Fig 12(b)**. The collected RTC acquired in Section 3.4 is utilized to convert the captured fluorescence signal from average pixel counts *I*_*mean*_ to imaged radiance *L*_*img*_, providing radiometric units of nW·cm^-2^·sr^-1^ for the captured fluorescence emission.

**Fig. 12:**
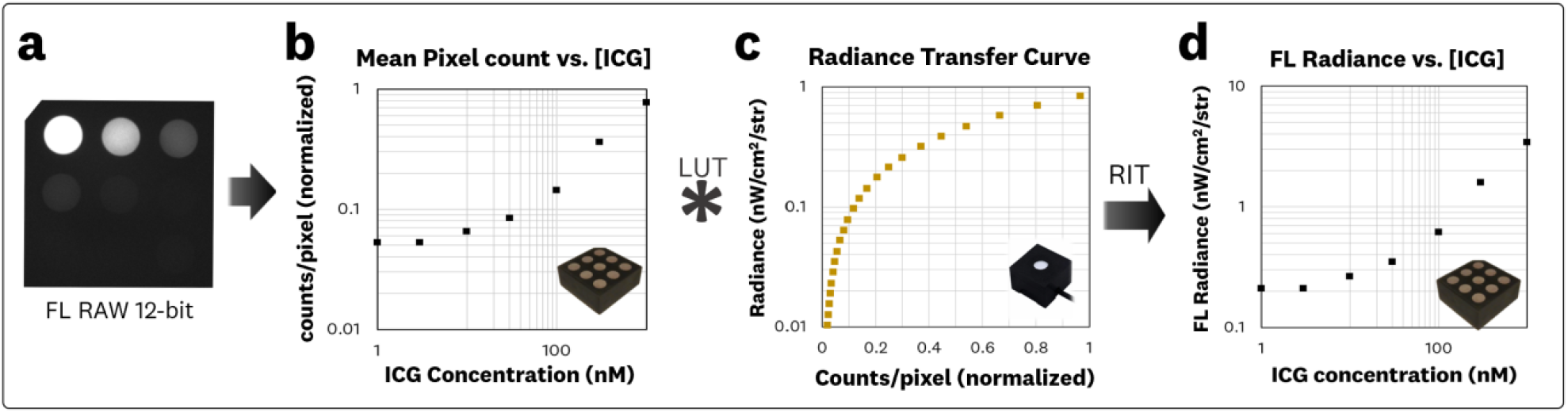
RIT transform of the acquired ICG concentration sensitivity target fluorescence image by utilizing the acquired RTC: **(a)** The resulting RAW 12-bit fluorescence image of the ICG concentration sensitivity target. **(b)** Plot of the average pixel counts *I*_*mean*_ (normalized to 12-bit depth) vs. ICG concentration for 5 mm ROIs on each of the nine wells. **(c)** The acquired RTC (Section 3.4) is utilized to perform the RIT on the acquired image which results in **(d)** the plot of imaged fluorescence radiance *L*_*img*_ *vs*. ICG concentrations.

#### 3.5.1 Assessing the RIT using varying image processing pipelines

To further explore the adequacy of the RTC and RIT to enable cross-system comparisons the captured RAW 12-bit fluorescence image of the ICG concentration target was post-processed utilizing an 8-bit down sampling and log10 compression that is re-mapped into a 12-bit image space, which are shown in **Fig 13(b)**. The corresponding mean pixel intensity (*I*_*mean*_) for each concentration well ROI for the images are plotted in Fig 13c, demonstrating significant differences in each dataset. These pipelines produced markedly different signal-to-background ratios with up to ∼306% disagreement (SBR_1000 nM_ = 15.3, 35.6, and 11.6 for the RAW 12-bit, 8-bit, and log-compressed images, respectively). Pipeline-specific RTCs [**Fig 13(d)**] for each image pipeline were used to perform the RIT which resulted in converging imaged radiance *L*_*img*_ values [**Fig 13(e)**]. The associated SBR_1000 nM_ values for the RIT imaged radiance were 17.4, 17.7, and 17.6 for the RAW 12-bit, 8-bit, and log10 12-bit images, respectively, showing an agreement within ∼2%. Significant improvements on reported metric agreement for the RIT images was also observed for the SNR and CNR metrics. For example, the SNR and CNR disagreements for the 1000 nM well decreased substantially, from approximately 495 % to 17 % and 504 % to 27 %, respectively, when comparing the unprocessed and RIT-processed images. In contrast, the CVR metric did not observe a significant improvement for the RIT images, showing reasonable agreement for the three differing imaging pipelines. The measured SBR, SNR, CNR, and CVR values of the wells for the unprocessed and RIT processed images are provided in **Table S1** and **Table S2** in the Supplementary Material.

**Fig. 13:**
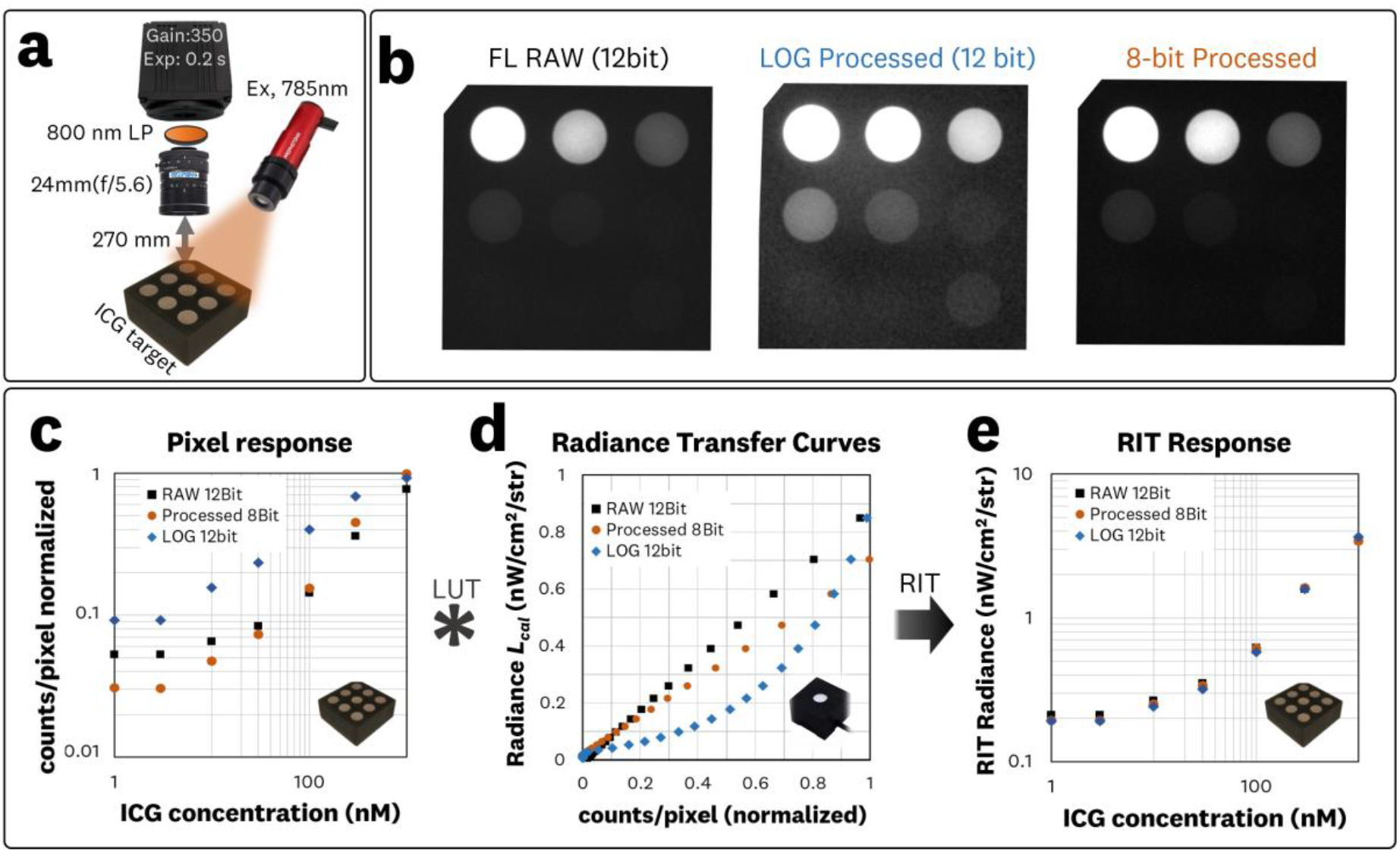
**(a)** Fluorescence imaging configuration for the ICG concentration sensitivity target **(b)** The RAW 12-bit image was post-processed through two pipelines: an 8-bit down-sampled image with baselining and a log10 compression that is re-mapped into a 12-bit image space. **(c)** Normalized mean pixel intensity (*I*_*mean*_) for 5-mm ROIs versus ICG concentration for the RAW, 8bit, and log10 images, showing disparity in the imaged responses. **(d)** Pipeline-specific RTCs were used to perform the RIT allowing for the conversion of the **(e)** pixel responses to imaged radiance (µW·cm^−2^·sr^−1^), resulting in converging fluorescence radiance vs. ICG concentration curves.

**Fig. 14:**
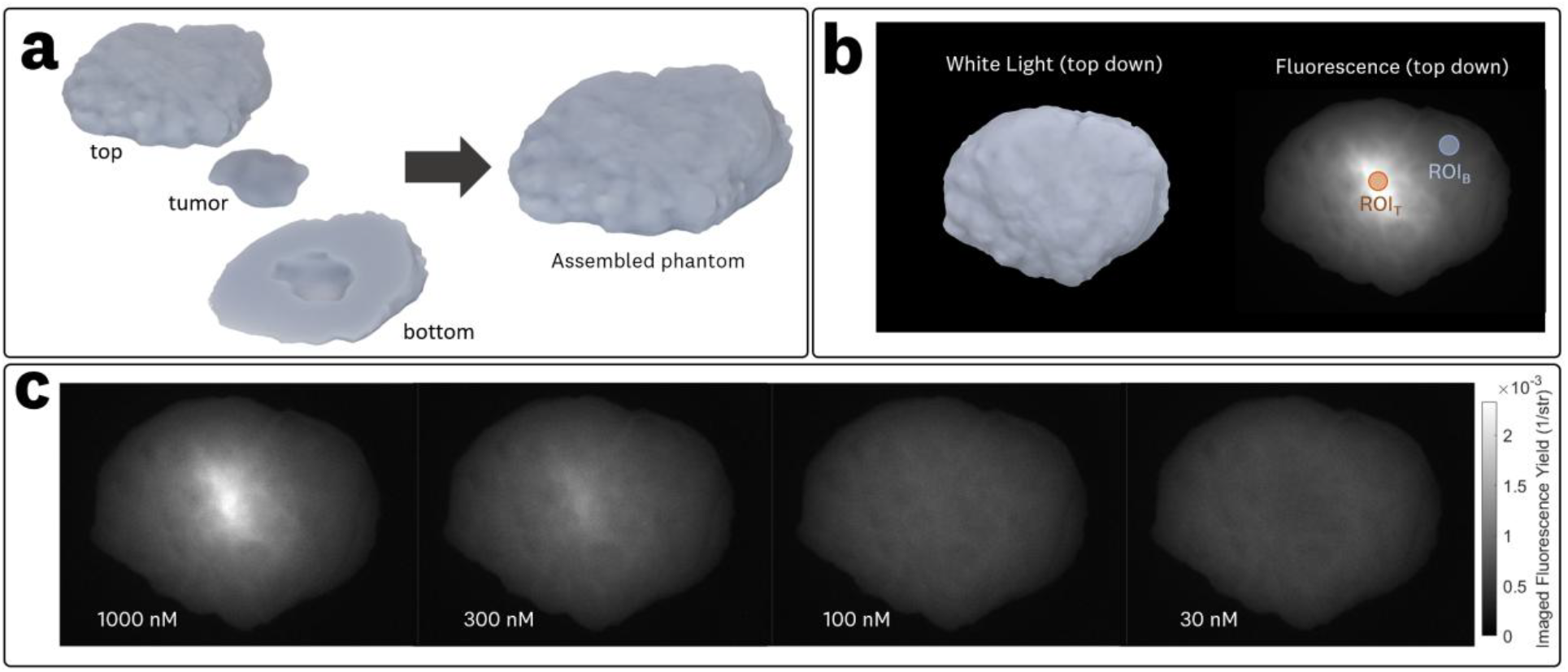
Breast lumpectomy phantom imaging (a) the solid 3D-printed phantom is composed of a top and bottom bulk material and a tumor inclusion with varying ICG concentrations. (b) Top-down white light and fluorescence images alongside the 5 mm ROIs for the tumor (ROI_T_) and background (ROI_B_). (c) The resulting FIT images for the 1000 nM, 300 nM, 100 nM, and 30 nM ICG tumor inclusions, providing SI-traceable units of str^-1^ and enabling the calculation of absolute imaging contrast metrics.

It is worth noting that because the excitation irradiance across the fluorescent target’s field of view was verified to be approximately uniform (>90% uniformity) in this experiment, contrast metrics derived from the RIT are effectively equivalent to those obtained from the FIT. However, under non-uniform illumination this equivalence does not hold, and the FIT with per-pixel normalization [**Equation (13)**] should be applied to correct for excitation-field inhomogeneities (see Section 2.6.2).

#### 3.5.2 Breast Lumpectomy Phantom Imaging

The imaging results of the breast lumpectomy phantom with tumor inclusions at ICG concentration of 30, 100, 300, 1000 nM are shown in **Fig 14**. The 5-mm ROIs used for the tumor and background calculation of mean and standard deviation are shown **Fig 14(b)**. The 12-bit RAW fluorescence images were converted from digital counts to imaged radiance (µW·cm^-2^·sr^-1^) using the RIT methodology with the RTC collected in Section 3.4. The excitation irradiance at the imaging plane was measured as 1.5 mW·cm^-2^, and used to normalize the fluorescence radiance and produce the FIT images shown in **Fig 14(c). Table S3** in the Supplementary Material reports the tumor and background ROI means and standard deviations in SI-traceable units of sr^-1^, representing the imaged aggregate fluorescence yield. The resulting SBR metrics were 3.7, 2.2, 1.4, and 1.1 for the 1000, 300, 100, and 30 nM inclusions, respectively. The resulting CNR metrics were 28.0, 13.4, 4.6, and 1.1 for the 1000, 300, 100, and 30 nM inclusions, respectively. The complete set of SBR, SNR, CNR, and CVR values computed from the 5-mm tumor and background ROIs for each inclusion configuration is provided in **Table S4** in the Supplementary Material.

## 4 Discussion

In this work, we established and assessed a radiometric framework for SI-traceable quantitative fluorescence imaging that combines a calibrated solid-state radiance emission target (RET) with three newly introduced methods: (1) the Radiance Transfer Curve (RTC), which captures sensor and image-processing nonlinearities across the usable signal range; (2) the Radiance Imaging Transform (RIT), which applies this calibration at the pixel level to map digital counts to SI-traceable radiance; and (3) the Fluorescence Imaging Transform (FIT), which performs pixel-wise excitation normalization of the RIT image to produce an aggregate fluorescence yield image. The RET’s Lambertian behavior was confirmed, with measured radiance remaining invariant to distance and aperture, supporting distance-invariant radiance transfer under fixed optical settings (Section 2.5, Appendix A). Imager responsivity *R*_*λ*_ was found to be stable across distance and aperture, and digital ROIs reproduced physical apertures, thereby enabling ROI and pixel-level radiance transfer. By sampling radiance across the detector’s dynamic range, the RTC extended this capability to the pixel level, allowing RIT conversion from counts to imaged radiance (µW·cm^−2^·sr^−1^). FIT further mapped images to aggregate fluorescence yield (sr^−1^), normalizing for excitation irradiance. Applied to an ICG concentration target, the RIT method harmonized outputs across the distinct image-processing pipelines (RAW, 8-bit, log10), yielding convergence of SBR/SNR/CNR. Finally, the breast lumpectomy phantom experiments demonstrated feasibility in anthropomorphic geometries and reported absolute contrast metrics from SI-traceable FIT. Finally, the breast lumpectomy phantom experiments demonstrated the feasibility of the approach in anthropomorphic geometries and produced absolute contrast metrics from SI-traceable FIT images, enabling true cross-system quantitative comparison.

### 4.1 Lambertian emission of the radiometric target

The radiance-versus-distance and radiance-versus-aperture experiments (Section 3.1, **Fig 7**) confirmed that the RET exhibited Lambertian emission for this on-axis orthogonal imaging. Across both experimental series, measured radiance showed no statistically significant dependence on distance or aperture, as supported by ANOVA results. Minor discrepancies at the largest working distance (400 mm) were attributed to reduced signal-to-noise, which in turn led to marginal outlier identification by robust statistical tests, but these did not alter the overall conclusion. Importantly, radiance remained constant within experimental uncertainty, providing empirical confirmation of Lambertian behavior. This invariance is essential for establishing SI-traceable radiance transfer. The RET thus provides a stable, calibrated source that enables the subsequent steps of the framework, including responsivity characterization, RTC generation, and RIT-based image conversion.

### 4.2 Conversion of radiance to camera response

The imager responsivity experiments (Section 3.2, **Fig 8**,**9**) further confirmed the Lambertian behavior of the RET and its suitability for imager response conversion and characterization. The results indicate that imager responsivity (*R*_*λ*_) was invariant, within experimental uncertainty, across the tested working distances and f-numbers, as expected from the Lambertian emission and *étendue* conservation described in Eqs. (3)–(8). This invariance validates the theoretical requirement for converting camera-specific responses into SI-traceable radiometric units. The *R*_*λ*_ departures observed at the smallest apertures (f/11) are most likely attributed to manufacturing tolerances of the aperture, which has been previously reported;^29^ for example, a 100 µm deviation for an f/11 aperture setting would be sufficient to account for the observed deviations. The differences in *R*_*λ*_ observed between lens/camera configurations are consistent with expected contributions from varying lens transmission, effective f-number, and acquisition parameters (exposure time, gain). Importantly, while a distance- and aperture-independent *R*_*λ*_ provides a local linear mapping from digital counts (*N*) to calibrated radiance (*L*_*cal*_) at a given operating point, it does not account for sensor or image-processing nonlinearities across the full dynamic range, motivating the development and evaluation of the RTC.

### 4.3 Physical vs. Digital apertures for radiance transfer

The physical-versus-digital aperture experiments (Section 3.3, **Fig 10**) showed that ROI masking produced imager response (*R*_*λ*_) values statistically indistinguishable from those obtained with matched physical apertures. This confirms that adjacent illuminated regions did not contribute appreciably to the ROI signal, enabling digital ROIs to substitute for physical apertures in responsivity estimation. Slightly lower *R*_*λ*_ values observed for the full 10 mm emission area reflect the RET’s pseudo top-hat profile, underscoring the need to account for spatial non-uniformity when extending calibrations from small ROIs to the full emission surface. Practically, this means that responsivity measurements utilizing the RET should use the entire emission surface as the ROI to minimize the impact of spatial non-uniformity. Overall, these results demonstrate that digital ROI selection is a valid approach for radiometric calibration and directly supports the ROI- and pixel-level radiance transfer required for RTC and RIT implementation.

### 4.4 Radiance Transfer Curve (RTC) for imager characterization

The RTC experiments (Section 3.4, **Fig 11**) expanded the characterization of imager response by sampling across the full dynamic range of calibrated RET radiances. Using a single responsivity value measured near 50% signal provided good agreement at moderate-to-high signals but produced significant deviations at lower signals, where sensor and processing nonlinearities become prominent. By acquiring images at 49 radiance levels spanning the detector’s noise floor to near saturation, the RTC established a direct mapping between mean pixel intensity and calibrated source radiance, thereby capturing the complete system response under fixed imaging parameters. As detailed in Appendix A, the RTC is invariant to source-to-aperture distance (i.e. working distance) because the Lambertian geometry preserves radiance transfer as long as other critical imaging parameters – aperture size (*A*_*apt*_), aperture-to-sensor distance (*d*_*sen*_), exposure time, gain, and optical components – are kept constant. Importantly, the RTC provides a comprehensive system characterization that extends beyond a single responsivity value, enabling precise pixel-level conversion to radiometric units and forming the foundation for the RIT and FIT methodologies.

### 4.5 Fluorescence Intensity Transform (RIT) and Fluorescence Imaging Transform (FIT)

The ICG concentration-sensitivity target experiments (Section 3.5; **Fig 12**,**13, Table S1**,**S2**) evaluated the RTC-enabled RIT across RAW, 8-bit down-sampled, and log10-compressed image pipelines. In device-count space, the different pipelines produced strikingly divergent responses, with contrast metrics such as SBR differing by several hundred percent depending on processing. After application of the RIT with pipeline-specific RTCs, these SBR discrepancies were reduced to within only a few percent, yielding closely aligned radiance–concentration curves. Comparable improvements were observed for SNR and CNR, while CVR remained less affected. These results highlight the sensitivity of conventional metrics to pipeline choices and demonstrate how the RIT and FIT provide a radiometrically consistent basis for quantitative fluorescence imaging across varying acquisition and image processing workflows.

I tis worth noting that the relatively larger discrepancies observed for the log10-compressed images likely originate from undersampling of low-radiance signal levels, where the logarithmic mapping disproportionately expands intensity differences; thus, further improvement may be achieved through finer sampling in the low-radiance regime for this processing pipeline. Additionally, these findings highlight the need for continued investigation into how image-processing pipelines influence variance-based performance metrics, particularly in the context of the improved reporting achieved through the RIT and FIT methodologies.

The breast lumpectomy phantom imaging (Section 3.5.2, **Fig 14**) demonstrated application of the RTC and RIT methodologies in anthropomorphic geometries. RAW fluorescence images were converted to RIT and subsequently normalized by excitation irradiance to generate FIT images with SI-traceable units of sr^-1^, enabling the calculation of absolute contrast metrics across tumor inclusions at varying ICG concentrations. While these experiments were primarily a feasibility evaluation, they demonstrated the capability of FIT to provide quantitative fluorescence imaging for standardized reporting in clinically relevant imaging. Importantly, the derived contrast metrics fell within ranges reported for clinical ICG imaging,^52–54^ indicating alignment with physiologically relevant conditions and supporting the potential use of RIT and FIT methodologies for standardized, quantitative reporting in clinical FGS and FGI studies.

### 4.6 Limitations and Future Work

The presented framework and evaluation are constrained by several underlying assumptions, which both support its development and delineate areas requiring further validation. First, the RET was characterized on-axis and used as a Lambertian calibrator; while the distance- and aperture-invariance results support this use, fluorescence from tissue can deviate from ideal Lambertian behavior due to surface effects, shallow scattering, and refractive-index mismatches. Off-axis RET characterization and angular emission tests are therefore warranted to confirm robustness across varied geometries and viewing angles. Second, RIT reports imaged radiance and FIT normalizes by the measured excitation irradiance at the sample plane; because excitation typically varies with working distance and illumination geometry, accurate irradiance measurement (and, when needed, pixel-wise uniformity correction)^47,55^ is essential for cross-system comparisons. Third, cross-system differences in excitation wavelength and bandwidth for a given fluorophore can alter the effective absorption cross-section and drive illumination-dependent nonlinearities (e.g., saturation, inner-filter effects, red-edge excitation shift,^56^ photobleaching), breaking the simple proportionality assumed by FIT; reporting the excitation spectrum/irradiance and, where feasible, applying spectral weighting and power-dependence checks will be important for harmonized comparisons. Fourth, the RET’s spectral band overlaps the ICG emission band for the configurations tested, but exact spectral matches will vary with fluorophore and filter sets; extending the approach with band-specific RTCs or spectral weighting would help address source–detector bandpass mismatches. Finally, while RIT/FIT operate at the pixel level, the current study used mostly planar, in-focus targets; future work should examine 3D surfaces within and beyond depth-of-field, where view angle and defocus could introduce small but systematic departures from the distance-invariant mapping assumed in Appendix A.

System integration and standardization also motivate further development. Many clinical imagers employ auto-gain, variable exposure, or high-dynamic-range stitching and may not persist complete acquisition metadata; some also include non-deterministic processing (e.g., denoising or machine-learning-based techniques). In such cases, a single RTC may be inadequate. A practical path is to (i) acquire families of RTCs across relevant settings (gain, exposure, f-number) or implement adapted radiometric self-calibration to interpolate between them;^42^ (ii) record and export the acquisition parameters and a unique RTC identifier alongside images as metadata; and (iii) include the excitation irradiance (and, when applicable, an illumination uniformity map). Because image-processing pipelines can alter noise statistics, future studies should analyze how this processing affects the standard deviations used in SNR, CNR, and CVR, and report metrics computed after RIT/FIT so that contrasts reflect radiometric – not pipeline-specific –differences. Lastly, a formal uncertainty budget for *L*_*cal*_, *L*_*img*_ and FIT is needed, including contributions from RET radiance calibration, aperture/entrance-pupil geometry, detector linearity, interpolation of the RTC LUT, excitation-irradiance measurement, and image non-uniformities.^29,31^ Establishing these uncertainties, extending RTC/RIT to additional spectral bands and fluorophores, and conducting multi-site evaluations on preclinical and FDA-cleared systems would help translate the framework into standardized, SI-traceable fluorescence reporting suitable for clinical studies.

## 5 Conclusion

Here we introduced a radiometric characterization framework for SI-traceable quantitative fluorescence imaging that combines a calibrated Lambertian emitter with three methods: the Radiance Transfer Curve (RTC), the Radiance Imaging Transform (RIT), and the Fluorescence Imaging Transform (FIT). We confirmed distance and aperture-invariant radiance transfer, showed that digital ROIs reproduce physical apertures, and demonstrated that a single responsivity does not capture low-signal and image processing induced nonlinearities, underscoring the need for an RTC that characterizes response across the full dynamic range of the device. Using acquired RTCs, the RIT mapped device-native counts to imaged radiance (µW·cm^−2^·sr^−1^) at the pixel level, with the FIT providing excitation-normalized aggregate fluorescence yield (sr^−1^) for the utilized emission and excitation bands. Applied to an ICG concentration target, the RIT/FIT reconciled large image processing pipeline-dependent disparities present in the device-specific responses. Breast-phantom imaging demonstrated feasibility in anthropomorphic geometries with absolute, SI-traceable contrast metrics reported. Together, these results indicate suitability of the presented framework to convert from device-specific images into radiometric and excitation-normalized outputs suitable for cross-system comparison and standardized reporting.

The feasibility results demonstrated here highlight the potential of the proposed framework to address the long-standing need for quantitative fluorescence imaging in FGI and FGS devices. Moving from feasibility to standardization will require additional development: establishing uncertainty budgets, validating performance across diverse optical geometries and illumination conditions, and accounting for inter-system differences in excitation spectra and emission collection. Equally important will be strategies to support routine use, including strategies to handle variable acquisition settings (families of RTCs or radiometric self-calibration) and exporting of acquisition parameters, RTCs, and measured excitation as image metadata. Additionally, broader evaluation across devices, fluorophores, and image processing pipelines is needed to establish robustness and generalizability, positioning the framework as a candidate for reproducible, quantitative fluorescence imaging that can be standardized across FGI and FGS systems.

## Appendix A: Derivation of the distance-invariant property of the Radiance Transfer Curve

Here, we derive RTC and radiance transfer distance-invariant property for a Lambertian extended source. As described in Section 2.4, the imaged flux *Φ*_*image*_ is given by **Equation (6)**. This radiant flux depends on the source radiance *L*_*src*_, source area *A*_*src*_, lens aperture *A*_*apt*_ and source-to-aperture distance *d*. However, in an imaging system, the source is not “smeared” over the entire sensor – the source area *A*_*src*_ is imaged onto a smaller area *A*_*img*_ on the sensor. The magnification *m* (approximated as *m* = *d*_*sen*_*/d* in a simple thin-lens system) resulting from imaging of the source gives:

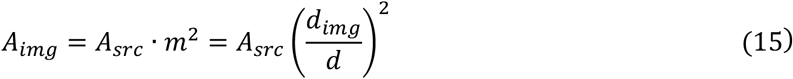

Next, we consider the irradiance (radiant flux per unit area) on the sensor, *E*_*img*_, which is given by:

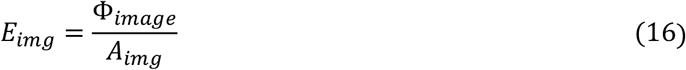

Substituting the expressions for *Φ*_*image*_ [**Equation (6)]** and *A*_*img*_ [**Equation (11)]** into **Equation (16)** we obtain:

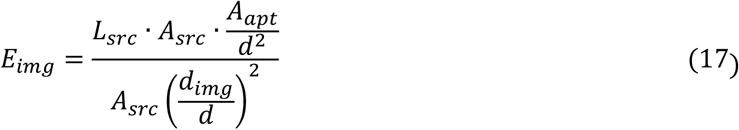

Where the *A*_*src*_ and *d* terms cancel out, simplifying to:

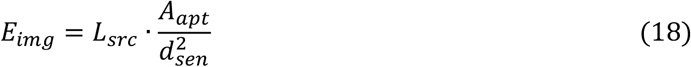

Such that the irradiance *E*_*img*_ associated with *A*_*img*_ is independent of the source-to-aperture distance *d* and instead depends only on the source radiance *L*_*src*_, lens aperture area *A*_*apt*_, and the (normally fixed) aperture-to-sensor distance *d*_*sen*_.

Next, we consider the camera conversion of the incident irradiance *E*_*img*_ into digital counts. We denote the irradiance conversion factor for a specified spectral band as *R*_*I,λ*_ (with units of counts/irradiance). The mean pixel intensity *I*_*mean*_, defined as the average count per pixel over the imaged area *A*_*img*_, is given by

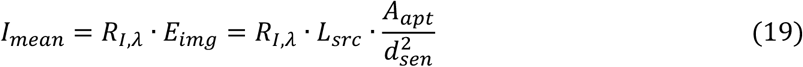

Since *R*_*I,λ*_, *A*_*apt*_, and *d*_*sen*_ are fixed for a given imaging configuration, we can define the radiance system responsivity for a given spectral band as *R*_*L,λ*_. It follows that we can simplify **Equation (19)** to

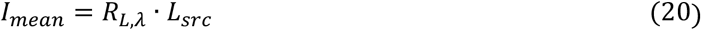

In other words, the mean pixel intensity *I*_*mean*_ is directly proportional to the source radiance *L*_*src*_ and independent of the source-to-aperture distance *d*.

In the context of the RTC calibration using the RET emission, where the radiance *L*_*cal*_ is a known SI-traceable radiance, we are essentially mapping *I*_*mean*_ ↔ *L*_*cal*_ over a range of *L*_*cal*_ values that span both the noise floor and saturation point of the imaging system response. This calibration results in a look-up-table of *R*_*L,λ*_ values that correlate the mean pixel intensities to SI-traceable radiance values. Because **Equation (20)** shows that for a Lambertian source the irradiance (and hence the pixel counts) is independent of *d*, once *A*_*apt*_ and *d*_*sen*_ are fixed, the RTC look-up table is valid for subsequent measurements regardless of the source distance. This invariance is a direct consequence of the conservation of *étendue* and the Lambertian source property, which together imply (1) the total radiant flux may change with *d*, but it is distributed over an image area that changes correspondingly and (2) the irradiance (radiant flux per unit area) on the sensor remains the same, so the digital count (or *I*_*mean*_ for a region-of-interest) remains directly tied to the source radiance. Although these results might appear counterintuitive, the data presented in Section 3.3.2 provide a clear visualization of these derivations, demonstrating that the pixel intensity of the imaged RET surface remains essentially constant across different source-to-aperture distances given fixed *A*_*apt*_ and *d*_*sen*_ parameters.

For a generalized derivation of the radiance distance invariance, readers are referred to Jannson (1980),^57^ which formulates a radiance-transfer framework suitable for advanced optical system modeling, incorporating diffraction, coherence, and imaging effects through propagation operators/kernels in the position–direction domain.

## Acknowledgments

This project was funded in part with federal funds from the National Institute of Biomedical Imaging and Bioengineering, National Institutes of Health, Department of Health and Human Services (Grant No. R43/44EB029804). The breast lumpectomy model utilized in this study was funded by the National Institutes of Health National Cancer Institute (NCI) contract 75N91021C00035. The authors also gratefully acknowledge Maritoni Litorja for her valuable guidance and discussions related to the radiance transfer and radiometric calibration aspects of this work.

## Code and Data Availability

The data used in this study are available from the corresponding author upon reasonable request.

## References

1. V. Ntziachristos, “FLUORESCENCE MOLECULAR IMAGING,” Annu. Rev. Biomed. Eng. 8(1), 1–33 (2006)[doi:10.1146/annurev.bioeng.8.061505.095831].

2. J. Rao, A. Dragulescu-Andrasi, and H. Yao, “Fluorescence imaging in vivo: recent advances,” Curr. Opin. Biotechnol. 18(1), 17–25 (2007) [doi:10.1016/j.copbio.2007.01.003].

3. R. K. Orosco, R. Y. Tsien, and Q. T. Nguyen, “Fluorescence Imaging in Surgery,” IEEE Rev. Biomed. Eng. 6, 178–187 (2013) [doi:10.1109/RBME.2013.2240294].

4. C. A. Combs, “Fluorescence Microscopy: A Concise Guide to Current Imaging Methods,” Curr. Protoc. Neurosci. 50(1) (2010) [doi:10.1002/0471142301.ns0201s50].

5. S. J. Sahl, S. W. Hell, and S. Jakobs, “Fluorescence nanoscopy in cell biology,” Nat. Rev. Mol. Cell Biol. 18(11), 685–701, Nature Publishing Group (2017) [doi:10.1038/nrm.2017.71].

6. F. Leblond et al., “Pre-clinical whole-body fluorescence imaging: Review of instruments, methods and applications,” J. Photochem. Photobiol. B 98(1), 77–94 (2010) [doi:10.1016/j.jphotobiol.2009.11.007].

7. A. Refaat et al., “In vivo fluorescence imaging: success in preclinical imaging paves the way for clinical applications,” J. Nanobiotechnology 20(1), 1–22, BioMed Central (2022) [doi:10.1186/s12951-022-01648-7].

8. N. Raison et al., “An evaluation of live porcine simulation training for robotic surgery,” J. Robot. Surg. 15(3), 429–434 (2021) [doi:10.1007/s11701-020-01113-3].

9. K. Joshi et al., “Pigs: Large Animal Preclinical Cancer Models,” World J. Oncol. 15(2), 149–168 (2024) [doi:10.14740/wjon1763].

10. I. L. Gitajn et al., “Evaluation of bone perfusion during open orthopedic surgery using quantitative dynamic contrast-enhanced fluorescence imaging,” Biomed. Opt. Express 11(11), 6458–6469, Optica Publishing Group (2020) [doi:10.1364/BOE.399587].

11. S. R. Thammineedi et al., “Fluorescence-guided cancer surgery—A new paradigm,” J. Surg. Oncol. 123(8), 1679–1698 (2021) [doi:10.1002/jso.26469].

12. M. D. Slooter et al., “Defining indocyanine green fluorescence to assess anastomotic perfusion during gastrointestinal surgery: systematic review,” BJS Open 5(2), Oxford Academic (2021) [doi:10.1093/bjsopen/zraa074].

13. P. A. Sutton et al., “Fluorescence-guided surgery: comprehensive review,” BJS Open 7(3), Oxford Academic (2023) [doi:10.1093/bjsopen/zrad049].

14. B. W. Pogue and E. L. Rosenthal, “Review of successful pathways for regulatory approvals in open-field fluorescence-guided surgery,” J. Biomed. Opt. 26(03) (2021) [doi:10.1117/1.JBO.26.3.030901].

15. M. Koch, P. Symvoulidis, and V. Ntziachristos, “Tackling standardization in fluorescence molecular imaging,” Nat. Photonics 12(9), 505–515 (2018) [doi:10.1038/s41566-018-0221-5].

16. B. W. Pogue et al., “Fluorescence-guided surgery and intervention — An AAPM emerging technology blue paper,” Med. Phys. 45(6), 2681–2688 (2018) [doi:10.1002/mp.12909].

17. B. Zhu and H. Jonathan, “A Review of Image Sensors Used in Near-Infrared and Shortwave Infrared Fluorescence Imaging,” Sensors 24(11), 3539 (2024) [doi:10.3390/s24113539].

18. M. I. Ochoa et al., “Assessment of open-field fluorescence guided surgery systems: implementing a standardized method for characterization and comparison,” J. Biomed. Opt. 28(9), 096007, SPIE (2023) [doi:10.1117/1.JBO.28.9.096007].

19. A. V. DSouza et al., “Review of fluorescence guided surgery systems: identification of key performance capabilities beyond indocyanine green imaging,” J. Biomed. Opt. 21(8) (2016) [doi:10.1117/1.JBO.21.8.080901].

20. P. DeRose and M. Litorja, “Comparison of calibration and standardization approaches for fluorescence guided imaging systems to benchtop fluorescence measurements in cellular systems,” in Design and Quality for Biomedical Technologies XI, R. Raghavachari, R. Liang, and T. J. Pfefer, Eds., p. 7, SPIE, San Francisco, United States (2018) [doi:10.1117/12.2301248].

21. U. Kanniyappan et al., “Performance test methods for near-infrared fluorescence imaging,” Med. Phys. 47(8), 3389–3401 (2020) [doi:10.1002/mp.14189].

22. A. J. Ruiz et al., “Indocyanine green matching phantom for fluorescence-guided surgery imaging system characterization and performance assessment,” J. Biomed. Opt. 25(05), 1 (2020) [doi:10.1117/1.JBO.25.5.056003].

23. P. A. Valdes et al., “Quantitative Wide-Field Imaging Techniques for Fluorescence Guided Neurosurgery,” Front. Surg. 6, |pnFrontiers (2019) [doi:10.3389/fsurg.2019.00031].

24. P. Holz and A. Brandenburg, “Calibration of systems for quantitative fluorescence analysis of thin layers,” Opt. Express 27(24), 34559 (2019) [doi:10.1364/OE.27.034559].

25. D. Gorpas et al., “Multi-Parametric Standardization of Fluorescence Imaging Systems Based on a Composite Phantom,” IEEE Trans. Biomed. Eng. 67(1), 185–192 (2020) [doi:10.1109/TBME.2019.2910733].

26. L. Hacker et al., “Criteria for the design of tissue-mimicking phantoms for the standardization of biophotonic instrumentation,” Nat. Biomed. Eng. 6(5), 541–558 (2022) [doi:10.1038/s41551-022-00890-6].

27. L. Wang, P. DeRose, and A. K. Gaigalas, “Assignment of the Number of Equivalent Reference Fluorophores to Dyed Microspheres,” J. Res. Natl. Inst. Stand. Technol. 121, 269 (2016) [doi:10.6028/jres.121.012].

28. C. Monte et al., “Linking fluorescence measurements to radiometric units,” Metrologia 43(2), S89–S93 (2006) [doi:10.1088/0026-1394/43/2/S18].

29. M. Litorja, “Conversion of imager-specific response to tissue phantom fluorescence into system of units-traceable units,” J. Biomed. Opt. 27(07) (2022) [doi:10.1117/1.JBO.27.7.074715].

30. M. Litorja et al., “Lambertian nature of tissue phantoms for use as calibrators in near infrared fluorescence imaging,” presented at SPIE BiOS, 10 March 2016, San Francisco, California, United States, 96960H [doi:10.1117/12.2216324].

31. B. Zhu et al., “Determining the Performance of Fluorescence Molecular Imaging Devices Using Traceable Working Standards With SI Units of Radiance,” IEEE Trans. Med. Imaging 35(3), 802–811 (2016) [doi:10.1109/TMI.2015.2496898].

32. M. Litorja, A. Urbas, and Y. Zong, “Radiometric calibration to consider in quantitative clinical fluorescence imaging measurements,” presented at SPIE BiOS, 4 March 2015, San Francisco, California, United States, 931114 [doi:10.1117/12.2086838].

33. B. Zhu et al., “Comparison of NIR Versus SWIR Fluorescence Image Device Performance Using Working Standards Calibrated With SI Units,” IEEE Trans. Med. Imaging 39(4), 944–951 (2020) [doi:10.1109/TMI.2019.2937760].

34. W. H. Steel, “Luminosity, Throughput, or Etendue?,” Appl. Opt. 13(4), 704 (1974) [doi:10.1364/AO.13.000704].

35. K. Niwa et al., “Quantitative Analysis of Bioluminescence Optical Signal,” Biosensors 13(2), 223, Multidisciplinary Digital Publishing Institute (2023) [doi:10.3390/bios13020223].

36. T. Enomoto et al., “Absolute bioluminescence imaging at the single-cell level with a light signal at the Attowatt level,” BioTechniques, Taylor & Francis (2018) [doi:10.2144/btn-2018-0043].

37. A. Sasaki and Y. Ohmiya, “Standardization of luminescence, fluorescence measurements, and light microscopy: Current situation and perspectives,” Biophys. Physicobiology 19(0), n/a (2022) [doi:10.2142/biophysico.bppb-v19.0037].

38. M. Yoshita et al., “Light-emitting-diode Lambertian light sources as low-radiant-flux standards applicable to quantitative luminescence-intensity imaging,” Rev. Sci. Instrum. 88(9), 093704 (2017) [doi:10.1063/1.5001733].

39. A. J. Ruiz et al., “Smartphone-based dual radiometric fluorescence and white-light imager for quantification of protoporphyrin IX in skin,” J. Biomed. Opt. 28(8), 086003, SPIE (2023) [doi:10.1117/1.JBO.28.8.086003].

40. J. R. Janesick, Photon Transfer, SPIE, 1000 20th Street, Bellingham, WA 98227–0010 USA (2007) [doi:10.1117/3.725073].

41. M. D. Grossberg and S. K. Nayar, “What Can Be Known about the Radiometric Response from Images?,” in Computer Vision — ECCV 2002 2353, A. Heyden et al., Eds., pp. 189–205, Springer Berlin Heidelberg, Berlin, Heidelberg (2002) [doi:10.1007/3-540-47979-1_13].

42. T. Mitsunaga and S. K. Nayar, “Radiometric self calibration,” in Proceedings. 1999 IEEE Computer Society Conference on Computer Vision and Pattern Recognition (Cat. No PR00149), pp. 374–380, IEEE Comput. Soc, Fort Collins, CO, USA (1999) [doi:10.1109/CVPR.1999.786966].

43. S. Lombardi and K. Nishino, “Radiometric Scene Decomposition: Scene Reflectance, Illumination, and Geometry from RGB-D Images,” in 2016 Fourth International Conference on 3D Vision (3DV), pp. 305–313, IEEE, Stanford, CA, USA (2016) [doi:10.1109/3DV.2016.39].

44. A. Litvinov and Y. Y. Schechner, “Radiometric framework for image mosaicking,” J. Opt. Soc. Am. A 22(5), 839 (2005) [doi:10.1364/JOSAA.22.000839].

45. E. Høg, “Photometric and Imaging Performance,” EAS Publ. Ser. 2, 27–37, EDP Sciences (2002) [doi:10.1051/eas:2002002].

46. N. Padmanabhan et al., “An Improved Photometric Calibration of the Sloan Digital Sky Survey Imaging Data,” Astrophys. J. 674(2), 1217–1233 (2008) [doi:10.1086/524677].

47. E. A. Mannoh et al., “Phantom for fluorescence uniformity and distortion assessment of near-infrared fluorescence guided surgery systems,” J. Biomed. Opt. 30(8), 086002, SPIE (2025) [doi:10.1117/1.JBO.30.8.086002].

48. J.-C. Pinoli, “The Logarithmic Image Processing Model: Connections with Human Brightness Perception and Contrast Estimators,” J. Math. Imaging Vis. 7(4), 341–358 (1997) [doi:10.1023/A:1008259212169].

49. N. Sadeghipour et al., “Prediction of optimal contrast times post-imaging agent administration to inform personalized fluorescence-guided surgery,” J. Biomed. Opt. 25(11) (2020) [doi:10.1117/1.JBO.25.11.116005].

50. E. P. M. LaRochelle et al., “3D-Printed Tumor Phantoms for Assessment of In Vivo Fluorescence Imaging Analysis Methods,” Mol. Imaging Biol. 25(1), 212–220 (2023) [doi:10.1007/s11307-022-01783-5].

51. A. J. Ruiz et al., “3D printing fluorescent material with tunable optical properties,” Sci. Rep. 11(1), 17135 (2021) [doi:10.1038/s41598-021-96496-0].

52. J. Feng et al., “Tumor-Background Ratio is an effective method to identify tumors and false-positive nodules in indocyanine-green navigation surgery for pediatric liver cancer,” Front. Pediatr. 10, 875688 (2022) [doi:10.3389/fped.2022.875688].

53. M. S. Kedrzycki et al., “The Impact of Temporal Variation in Indocyanine Green Administration on Tumor Identification During Fluorescence Guided Breast Surgery,” Ann. Surg. Oncol. 28(10), 5617–5625 (2021) [doi:10.1245/s10434-021-10503-2].

54. C. F. Pop et al., “Fluorescence imaging for real-time detection of breast cancer tumors using IV injection of indocyanine green with non-conventional imaging: a systematic review of preclinical and clinical studies of perioperative imaging technologies,” Breast Cancer Res. Treat. 204(3), 429–442 (2024) [doi:10.1007/s10549-023-07199-1].

55. W. D. Niles and F. S. Cohen, “Radiometric calibration of a video fluorescence microscope for the quantitative imaging of resonance energy transfer,” Rev. Sci. Instrum. 66(6), 3527–3536, AIP Publishing (1995) [doi:10.1063/1.1145465].

56. A. J. Ruiz et al., “Indocyanine green excitation-emission matrix characterization: spectral shifts and application-specific spectra.”

57. T. Jannson, “Radiance transfer function,” J. Opt. Soc. Am. 70(12), 1544 (1980) [doi:10.1364/JOSA.70.001544].

