## Supplementary Material for "Quantitative Fluorescence Imaging with SI-Traceable Radiance: Radiometric Fluorescence Characterization with a Calibrated Solid-State Emitter"

**Table S1**: Calculated imaging contrast metrics for the unprocessed images of the ICG reference concentration sensitivity (RCS) target. The calculated metrics were signal-to-background ratio (SBR), signal-to-noise ratio (SNR), contrast-to-noise ratio (CNR), and contrast-to-variability ratio (CVR) computed from 5-mm ROIs on each well for all three image-processing pipelines: 12-bit RAW, 8-bit down-sampled with baselining, and log10-compressed with 12-bit remapping.

|  | | **RCS ICG Target Well (nM)** | | | | | | |
| --- | --- | --- | --- | --- | --- | --- | --- | --- |
| **Contrast Metric** | **Image** | **1000** | **300** | **100** | **30** | **10** | **3** | **1** |
| **SBR** | 12bit RAW | 15.3 | 7.1 | 2.8 | 1.7 | 1.3 | 1.0 | 1.0 |
|  | 8bit | 35.6 | 16.3 | 5.6 | 2.6 | 1.7 | 1.1 | 1.1 |
|  | log10 | 11.6 | 8.6 | 5.0 | 3.0 | 2.0 | 1.2 | 1.1 |
| **SNR** | 12bit RAW | 113.4 | 52.9 | 21.0 | 12.3 | 9.5 | 7.7 | 7.7 |
|  | 8bit | 105.0 | 48.1 | 16.4 | 7.8 | 5.1 | 3.3 | 3.2 |
|  | log10 | 22.9 | 17.1 | 10.0 | 5.8 | 3.9 | 2.3 | 2.3 |
| **CNR** | 12bit | 105.9 | 45.4 | 13.6 | 4.9 | 2.1 | 0.3 | 0.3 |
|  | 8bit | 102.0 | 45.1 | 13.5 | 4.8 | 2.1 | 0.3 | 0.3 |
|  | log | 21.0 | 15.1 | 8.0 | 3.9 | 1.9 | 0.3 | 0.3 |
| **CVR** | 12bit RAW | 18.9 | 14.1 | 6.7 | 3.0 | 1.4 | 0.2 | 0.2 |
|  | 8bit | 37.6 | 14.1 | 6.7 | 3.0 | 1.4 | 0.2 | 0.2 |
|  | log10 | 19.6 | 13.8 | 6.7 | 3.0 | 1.4 | 0.2 | 0.2 |

**Table S2**: Calculated imaging contrast metrics for the RIT-processed images of the ICG RCS target after applying pipeline-specific radiometric transfer curves (RTCs) to obtain SI-traceable imaged radiance. Metrics (SBR, SNR, CNR, and CVR) were computed from the same 5-mm ROIs used in Table S1, across the image processing pipelines: 12-bit RAW, 8-bit down-sampled, and log10-compressed with 12-bit remapping.

|  | | **RCS ICG Target Well (nM)** | | | | | | |
| --- | --- | --- | --- | --- | --- | --- | --- | --- |
| **Contrast Metric** | **Image** | **1000** | **300** | **100** | **30** | **10** | **3** | **1** |
| **SBR** | 12bit RAW | 17.4 | 8.1 | 3.1 | 1.8 | 1.3 | 1.0 | 1.0 |
|  | 8bit | 17.7 | 8.1 | 3.1 | 1.8 | 1.3 | 1.0 | 1.0 |
|  | log10 | 17.6 | 8.2 | 3.2 | 1.8 | 1.4 | 1.1 | 1.1 |
| **SNR** | 12bit RAW | 110.6 | 51.5 | 20.0 | 11.2 | 8.5 | 6.7 | 6.7 |
|  | 8bit | 110.4 | 50.3 | 19.5 | 11.0 | 8.3 | 6.5 | 6.5 |
|  | log10 | 94.1 | 43.9 | 17.1 | 9.6 | 7.3 | 5.7 | 5.7 |
| **CNR** | 12bit | 104.3 | 45.1 | 13.6 | 4.9 | 2.1 | 0.3 | 0.3 |
|  | 8bit | 104.2 | 44.1 | 13.3 | 4.7 | 2.1 | 0.3 | 0.3 |
|  | log | 81.7 | 35.4 | 10.8 | 3.9 | 1.8 | 0.3 | 0.3 |
| **CVR** | 12bit RAW | 19.2 | 14.1 | 6.8 | 3.0 | 1.4 | 0.2 | 0.2 |
|  | 8bit | 27.3 | 14.0 | 6.7 | 2.9 | 1.4 | 0.2 | 0.2 |
|  | log10 | 19.8 | 13.7 | 6.4 | 2.7 | 1.3 | 0.2 | 0.2 |

**Table S4**: Breast lumpectomy phantom measured tumor and background ROI means and standard deviations from the FIT images, reported in SI-traceable aggregate fluorescence yield units (sr⁻¹). Data are shown for each tumor inclusion configuration of 1000, 300, 100, and 30 nM ICG. The reported units are 10^-3^∙sr^-1^.

| **Inclusion**  **Concentration** | **Tumor ROI** | | **Background ROI** | |
| --- | --- | --- | --- | --- |
|  | **Mean** | **Std** | **Mean** | **Std** |
| **1000nM** | 2.14 | 0.14 | 0.58 | 0.06 |
| **300nM** | 1.20 | 0.07 | 0.55 | 0.05 |
| **100nM** | 0.73 | 0.05 | 0.51 | 0.05 |
| **30 nM** | 0.58 | 0.04 | 0.53 | 0.04 |

**Table S4**: Breast lumpectomy phantom contrast metrics SBR, SNR, CNR, and CVR derived from 5-mm tumor and background ROIs of the FIT image for each tumor inclusion configuration of 1000, 300, 100, and 30 nM ICG.

| **Contrast Metric** | **Breast Tumor ICG concentration** | | | |
| --- | --- | --- | --- | --- |
|  | **1000nm** | **300nm** | **100nm** | **30nm** |
| **SBR** | 3.7 | 2.2 | 1.4 | 1.1 |
| **SNR** | 38.6 | 24.7 | 15.7 | 13.0 |
| **CNR** | 28.0 | 13.4 | 4.6 | 1.1 |
| **CVR** | 10.6 | 7.3 | 3.3 | 0.8 |
